# Engagement of spinal- vs prefrontal-projecting locus coeruleus neurons parses analgesia from aversion & anxiety in rats

**DOI:** 10.1101/156307

**Authors:** Stefan Hirschberg, Yong Li, Andrew Randall, Eric J Kremer, Anthony E Pickering

## Abstract

The locus coeruleus (LC) projects throughout the brain and spinal cord and is the major source of central noradrenaline. It remains unclear whether the LC acts functionally as a single global effector or as discrete modules. Specifically, while spinal-projections from LC neurons can exert analgesic actions, it is not known whether they can act independently of ascending LC projections. Using viral vectors taken up at axon terminals, we expressed chemogenetic actuators selectively in LC neurons with spinal (**LC^:SC^**) or prefrontal cortex (**LC^:PFC^**) projections. Activation of the **LC^:SC^** module produced robust, lateralised analgesia while activation of **LC^:PFC^** produced aversion. In a neuropathic pain model, **LC^:SC^** activation reduced hind-limb sensitization and induced conditioned place preference. By contrast, activation of **LC^:PFC^** exacerbated spontaneous pain, produced aversion and increased anxiety-like behaviour. This independent, contrasting modulation of pain-related behaviours mediated by distinct noradrenergic modules provides evidence for a discrete functional organisation of the LC.

## INTRODUCTION

Neuropathic pain has a population prevalence of 7-8% (Bouhassira et al., 2008, Torrance et al., 2006) and remains difficult to treat (Finnerup et al., 2015) mandating research to improve therapies (von Hehn et al., 2012, Skolnick and Volkow, 2016). Many of the frontline treatments for neuropathic pain modulate spinal noradrenaline, either directly such as noradrenaline re-uptake inhibitors or as part of a common effector pathway such as gabapentinoids and opioids (Nakajima et al., 2012, Hayashida et al., 2007, Hayashida et al., 2008, Jasmin et al., 2003). A functional deficit of noradrenergic control is associated with the development of sensitisation in neuropathic pain models (Hughes et al., 2013, Hughes et al., 2015, De Felice et al., 2011, Xu et al., 1999). However, current systemic pharmacological interventions to correct this imbalance commonly produce CNS side-effects such as anxiety, sleep disturbance, mood shifts and confusion, limiting their therapeutic utility (Finnerup et al., 2015).

The locus coeruleus (LC) is the principal noradrenergic nucleus in the CNS (Dahlstroem and Fuxe, 1965) and is the main source of noradrenergic innervation to the spinal dorsal horn, forming part of a well-described analgesic circuit (Millan, 2002, Pertovaara, 2006, Howorth et al., 2009a, Bruinstroop et al., 2012). The LC also projects throughout almost the entire neuraxis and plays a pivotal role in diverse behaviours such as learning and memory (Takeuchi et al., 2016, Martins and Froemke, 2015), strategic behaviour (Tervo et al., 2014), motivation (Varazzani et al., 2015), and arousal (Sara and Bouret, 2012, Carter et al., 2010, Vazey and Aston-Jones, 2014). Furthermore, LC-derived noradrenaline is also causally linked to aversive states like stress, anxiety and depression (McCall et al., 2015, Alba-Delgado et al., 2013). Thus, drug treatments for chronic pain that globally enhance noradrenaline levels or mimic its action are likely to cause adverse effects by acting on LC neuronal circuits that are not directly involved in pain control.

The LC has been characterised as a homogeneous cluster of neurons that provide a uniform global signal as part of a central arousal system (Fuxe et al., 2010). There is, however, accumulating evidence for specialization of the LC neurons into subsets on the basis of: 1) anatomical projections (Loughlin et al., 1986, Kebschull et al., 2016, Chandler et al., 2014); 2) electrical properties (Chandler et al., 2014); and 3) co-transmitter content (Simpson and Lin, 2007). In accord with this organisational principle we have demonstrated that distinct populations of LC noradrenergic neurons innervate the spinal cord (**LC^:SC^**) and prefrontal cortex (**LC^:PFC^**) based on anatomy and electrophysiology *in vitro* and *in vivo* (Li et al., 2016). Here we selectively target LC neurons from their projections to the spinal cord or prefrontal cortex using the axon terminal uptake and retrograde transport of canine adenovirus type 2 (CAV-2) vectors (Junyent and Kremer, 2015, Li et al., 2016). This CAV-2 vector expresses a chemogenetic actuator in noradrenergic neurons specifically, to allow activation of **LC^:SC^** or **LC^:PFC^** neurons to test whether the beneficial noradrenergic analgesia and stress-like adverse effects seen in pain states are mediated by distinct subpopulations of LC neurons and are therefore dissociable.

## Results

### Targeting and activation of noradrenergic LC neurons with PSAM

To enable selective activation of LC neurons, the excitatory chemogenetic receptor PSAM (*Pharmaco-Selective Actuator Module, PSAM ^L141F,Y115F:5HT3HC^ (Magnus et al., 2011)*) was expressed in noradrenergic neurons (**Figure 1**A-B) using viral vectors with the cell-type specific synthetic promotor PRSx8 (PRS) (Hwang et al., 2001, Lonergan et al., 2005, Howorth et al., 2009a). Efficient and selective targeting of LC neurons was observed after direct injection into the dorsal pons of a lentiviral vector harbouring the PRS-EGFP-2A-PSAM expression cassette. After transduction, EGFP immunofluorescence was detected in 690±38 neurons per LC (N=5) of which 98±0.4% co-expressed dopamine β-hydroxylase (DBH) confirming their identity as noradrenergic (**Figure 1**B). To enable post-hoc immuno-localisation, PSAM was tagged with a C-terminal HA. Subsequently, PSAM-HA was detected in the membranes of the LC neurons (**Figure** 1C, 82/112 transduced neurons examined from 3 rats). The functionality of the PSAM-HA was verified by Ca^2+^-imaging in cell lines (**Figure S 1**).

**Figure 1.**
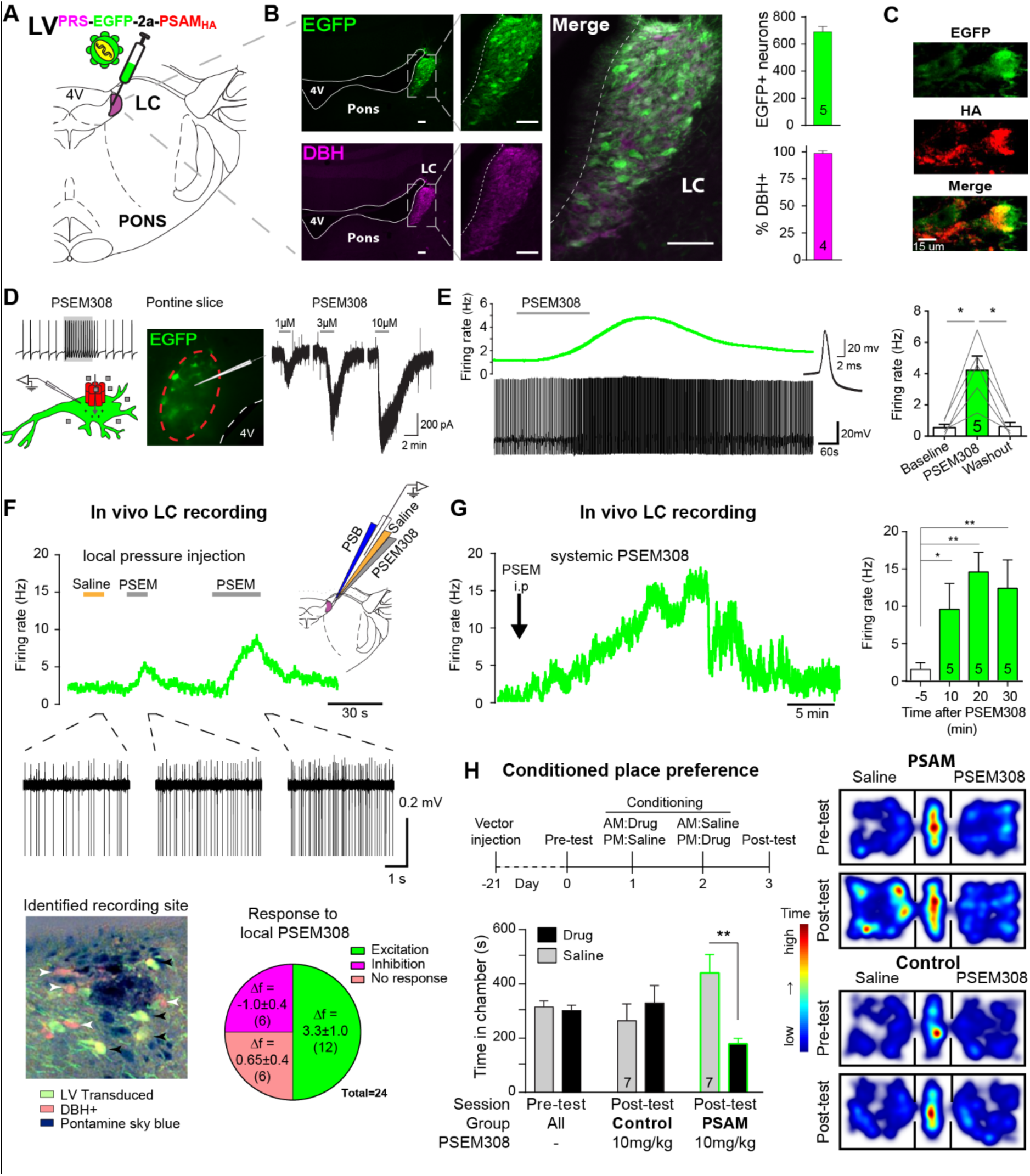
PSAM mediated chemogenetic activation of LC neurons in vivo. (A) Strategy using direct stereotaxic injection of lentiviral vector to express the excitatory ionophore PSAM in noradrenergic LC neurons. (B) Selective transduction of LC demonstrated by immunohistochemistry (IHC) for EGFP and dopamine β-hydroxylase (DBH) with 690 EGFP+ neurons per LC of which 98% were DBH+ (scale bar 100 μm). (C) PSAM expression was demonstrated using IHC for the HA tag. (scale bar 15 μm) (D) Schematic of PSEM308-mediated excitation of transduced neurons expressing PSAM. Patch clamp recordings from EGFP+ LC neurons in acute pontine brain slices. Perfusion of PSEM308 evoked concentration-dependent inward currents. (E) PSEM308 (3μM) increased rate of firing of transduced LC neurons. Inset shows 16 overlaid action potentials. Group data shows increase in firing produced by PSEM308 (3μM). (F) Extracellular recordings from LC neurons in anaesthetised rats using multi-barrel recording electrodes allowing local pressure ejection of PSEM308/saline/pontamine sky blue (PSB). Traces show graded excitation of an identified LC neuron by PSEM308. Recording sites were subsequently histologically identified by the PSB staining within the LC (transduced cells identified by IHC for EGFP (black arrowheads) and DBH (white arrowheads). The response to local PSEM308 was categorised as excitation or inhibition if it changed firing rate by more than 3 SD from the baseline rate. Application of PSEM308 produced an excitation in 50% of LC neurons. We found a second group of neurons that showed no response, presumably as they were not transduced. A third group showed an inhibition of spontaneous firing in response to local PSEM308 application. (G) Kinetics of the excitatory response to systemic PSEM308 administration (10 mg/kg i. p). (H) Timeline of conditioned place aversion protocol to assess influence of chemogenetic activation of LC neurons on behaviour. In PSAM expressing rats, PSEM308 (10 mg/kg) caused conditioned place aversion but had no effect on control animals. Representative heat maps of rat position in the pre-test and post-test after PSEM308 with bilateral LC transduction with LV^PRS-EGFP-2A-PSAM HA^ or control LV^PRS-EGFP^ All data analysed with repeated measures ANOVA (one or two way as appropriate) with Bonferroni’s post hoc testing (* P<0.05, ** P<0.01). (See also Figure S 1 and Figure S 2)

Patch clamp recordings from transduced LC neurons in acute pontine slices showed that the designer-ligand PSEM308 (*pharmaco-selective effector molecule)* produced inward currents and a reversible excitation. PSEM308 evoked concentration-dependent inward currents of up to 800pA in voltage clamp (V_hold_ -60 mV, 1-10 μM, **Figure 1**D). Current-clamp recordings of transduced LC neurons showed an 8-fold increase in firing frequency (0.54±0.21 Hz to 4.22±0.90 Hz, **Figure 1**E) in response to PSEM308 (3 μM). No response to PSEM308 (doses up to 10 μM) was detected in recordings from nontransduced LC neurons (n=6). We found no difference in intrinsic electrical properties between transduced and non-transduced LC neurons (**Table S 2**) suggesting that expression of PSAM is well tolerated. Together, these findings indicate that PSAM is functionally expressed in LC neurons and its activation by its ligand is sufficient to robustly increase action potential discharge.

### Chemogenetic excitation of the LC in vivo

Extracellular recordings with multi-barrelled electrodes were used to determine whether PSEM308 application could excite LC neurons *in vivo* (after direct LC transduction with LV^PRS-EGFP-2A-PSAM^ (**Figure 1**F)). In PSAM expressing rats (N=8), local pressure application of PSEM308, but not saline, caused a time-locked and reproducible excitation in 12 out of the 24 recorded units (**Figure 1**F), increasing their firing to 234% of baseline (2.63±0.92 Hz to 6.15±1.47 Hz). The remaining cells did not respond (n=6) or showed a small reduction of their firing frequency (-34.6%, 2.83±1.21 to 1.85±0.86 Hz, n=6) in response to local PSEM308 (**Figure 1**F). These inhibited units, which were in some cases simultaneously recorded along with LC units whose activity was increased (**Figure S 2**), likely reflect LC neurons that are not themselves transduced but are inhibited by noradrenaline released from surrounding transduced and hence PSEM308-activated LC neurons (as previously hypothesised by Vazey and Aston-Jones (2014)) These data are in line with pontamine sky blue labelling of the recording sites, which were always surrounded by DBH expressing neurons of which many, but not all, were also EGFP+ (**Figure 1**F). Local application of PSEM308 did not alter baseline firing frequency in control LC neurons (2.99±1.55 to 3.15±1.58 Hz, NS, n=7 neurons from 3 non-transduced rats).

The effect of systemic application of PSEM308 was tested on transduced LC units that showed an excitatory response to local PSEM308 (n=5, **Figure 1**G). Intraperitoneal (i.p) injection of PSEM308 (10 mg/kg, same dose used throughout the rest of the study unless otherwise stated) increased discharge frequency 10-fold from 1.47±0.93 to 14.76±2.8 Hz after 20 min. LC neurons doubled their firing frequency after 5.9 ± 3.9 min and cells reached their maximum discharge frequency after 15.9 ± 6.5 min (**Figure 1**G, median 15.8 Hz, range 10.5 to 27.2 Hz). The evoked excitation outlasted the time of experimental recording (30 min) in four out of five neurons.

### Chemogenetic activation of the LC alters behaviour

High tonic LC discharge frequencies (5-10Hz) are acutely aversive (McCall et al., 2015). Therefore, we employed a Pavlovian-conditioned place preference (CPP) paradigm to assay whether there was a behavioural consequence to chemogenetic activation of LC neurons. CPP is a voluntary choice task using the innate behaviour of animals to divide their time between environments according to their affective valence. Conditioning with PSEM308 induced avoidance of the drug-paired environment in animals that bilaterally expressed PSAM in LC neurons (**Figure 1**H). Such avoidance was not seen with a lower dose of PSEM308 (5 mg/kg), or in animals transduced with a control vector (LV^PRS-EGFP^) and administered PSEM308 (10 mg/kg).

### Activation of LC^:SC^ but not LC^:PFC^ is analgesic

Based on previous studies we predicted that **LC^:SC^** neurons would play a role in anti-nociception (Jones and Gebhart, 1986, Howorth et al., 2009b) and hypothesised that the forebrain projecting LC neurons might play a different – potentially pro-nociceptive role (Hickey et al., 2014). To target noradrenergic neurons according to their projection territory, we inserted a PRS-EGFP-2A-PSAM expression cassette into a CAV-2 vector to target LC neurons (Figure 2A) from their axonal projections (Li et al., 2016, Soudais et al., 2001).

**Figure 2.**
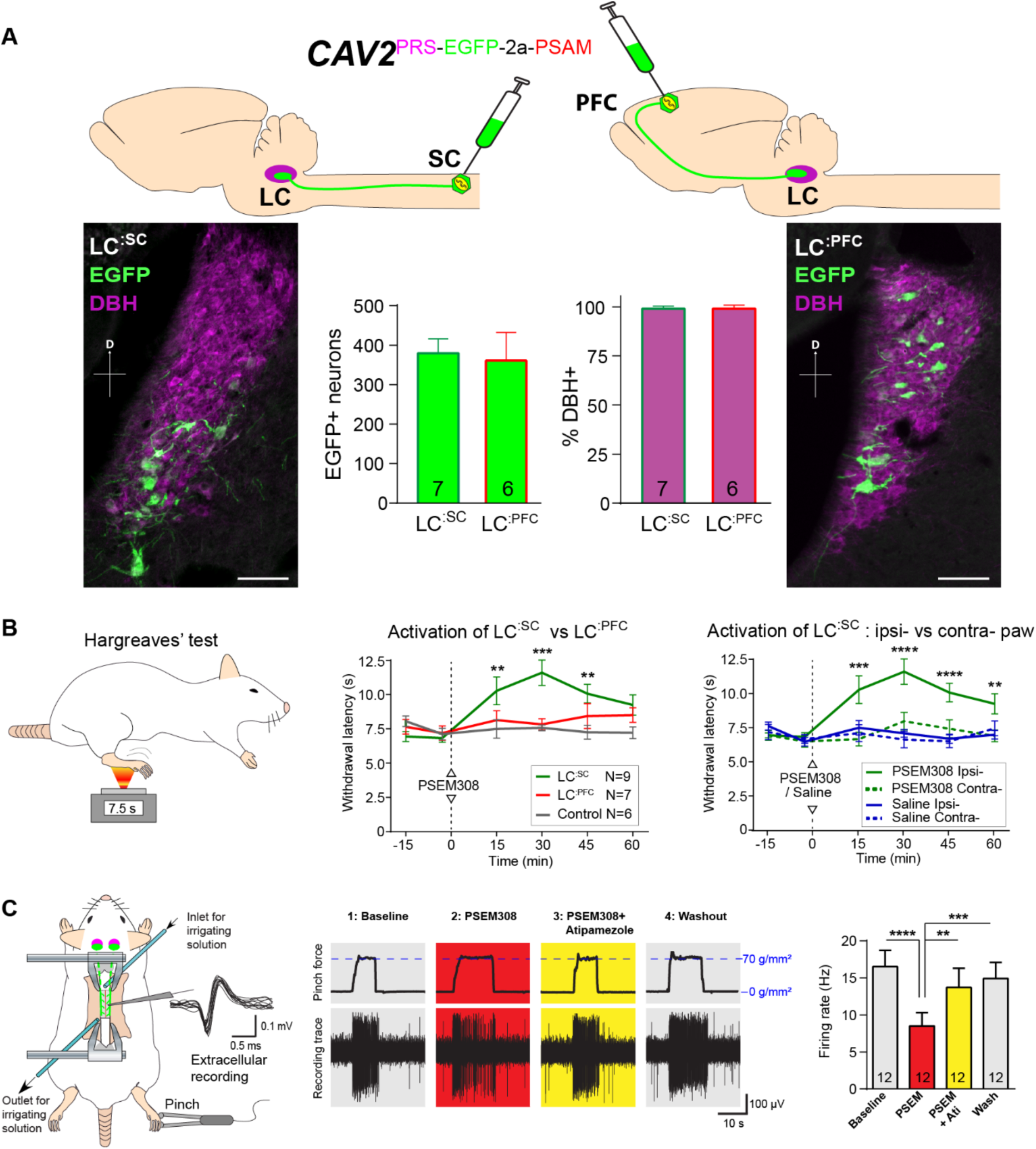
Activation of descending **LC^:SC^** but not ascending LC^:PFC^ is analgesic. (A) Retrograde transduction strategy with canine adenoviral vectors (CAV2) to target noradrenergic LC neurons with projections to the spinal cord (**LC^:SC^**) or prefrontal cortex (**LC^:PFC^**). Similar numbers of LC neurons were transduced by injections in lumbar spinal cord (L3-4, 380) and prefrontal cortex (361). In both cases >99% of neurons were DBH+. (scale bar 100 μm). (B) The Hargreaves’ test (radiant heat) was used to measure hind-paw withdrawal latency. PSEM308 activation of **LC^:SC^** but not **LC^:PFC^** caused a robust analgesic effect (increase in withdrawal latency). The **LC^:SC^** analgesic effect was only seen in the ipsilateral hind paw (same side as spinal CAV2 injection). (C) Spinal extracellular recording from wide dynamic range neurons. Calibrated forceps applied noxious pinch to hindpaw during spinal drug application. In **LC^:SC^** transduced rats, spinal superfusion of PSEM308 (10 μM) caused a significant attenuation of the response to noxious pinch which was prevented by Atipamezole (10 μM) indicating a spinal α2-adrenoceptor mechanism. All data analysed with 2-way repeated measures ANOVA with Bonferroni’s post hoc testing (** P<0.01, ***P<0.001, ****P<0.0001). (See also Figure S 3)

Bilateral injection of CAV2^PRS-EGFP-2A-PSAM^ into the dorsal horn of the lumbar spinal cord transduced 380±37 neurons in each LC (N=7, **Figure 2**A). These **LC^:SC^** neurons were located in the ventral aspect of the nucleus. In a second set of animals, CAV2^PRS-EGFP-2A-PSAM^ bilaterally injected into the PFC transduced similar numbers of neurons (361±72 per LC, N=6). However, these **LC^:PFC^** neurons were scattered throughout the LC with a more dorsal location than the **LC^:SC^** neurons (representing different groups of neurons as previously noted (Bruinstroop et al., 2012, Howorth et al., 2009a, Li et al., 2016)). In both cases, the transduced **LC^:SC^** and **LC^:PFC^** neurons were positive for DBH (99.1±0.5% and 99.1±0.7% respectively) indicating efficient and specific retrograde transduction of the noradrenergic neurons (**Figure 2**A).

We used the Hargreaves’ test to measure hind paw thermal withdrawal latency as an assay of nociception. Chemogenetic activation of **LC^:SC^** neurons (PSEM308 i.p, N=7 rats) significantly increased thermal withdrawal latency to radiant heat as compared to naïve rats (N=6) (30min post i.p., 11.6±0.9 s **LC^:SC^** vs 7.6±0.2 s naïve (**Figure 2**B)). This time course of analgesic effect between 15 and 45 min post-dosing is consistent with the kinetics of LC neuron activation after i.p PSEM308 injection (**Figure 1**G). Moreover, this **LC^:SC^** analgesia was only seen in the hind-limb ipsilateral to the vector injection in the spinal cord (**Figure 2**B), indicating an unexpected lateralised organisation of the **LC^:SC^** projection. By contrast, activation of **LC^:PFC^** neurons had no effect on the thermal withdrawal latency. This indicates a functional dichotomy in the actions of the ascending and descending LC projections on acute pain.

To investigate the mechanism of **LC^:SC^** chemogenetic analgesia, extracellular recordings were made from wide dynamic range (WDR) neurons in the spinal dorsal horn (n=14 from 6 rats, mean depth 415 μm, range 200 to 600 μm) in a preparation that allows the application of drugs directly onto the spinal cord in the irrigating solution (**Figure 2**C, (Funai et al., 2014, Furue et al., 1999)). WDR neurons encode stimulus intensity, responding to both innocuous and noxious stimuli (Price and Dubner, 1977). The average spontaneous discharge frequency of WDR was 1.86 ± 0.60 Hz, that was increased to 16.52 ± 2.19 Hz by pinch stimulation of the hind paw. In **LC^:SC^** transduced animals, perfusion of PSEM308 (10 μM) inhibited the pinch-evoked discharge two-fold to 8.5 ± 1.8 Hz (12/14 WDR neurons, **Figure 2**C). However, both spontaneous firing and innocuous, brush-evoked discharge were unaffected by PSEM308 (**Figure S 3**). These data demonstrate that local activation of terminals of **LC^:SC^** neurons in the spinal cord causes a specific inhibition of nociceptive inputs to WDR neurons.

To address the question of whether the **LC^:SC^** inhibition of the pinch response was caused by spinal release of noradrenaline, Atipamezole (10 μM, α2-adrenoceptor antagonist) was co-applied with PSEM308 (10 μM). The presence of atipamezole prevented the **LC^:SC^** mediated-inhibition of the pinch response (**Figure 2**C). Atipamezole alone had no effect on spontaneous discharge, evoked discharge after pinch or brush indicating there is little ongoing spinal release of noradrenaline under anaesthesia (paired t-test p>0.05, n=5 WDR neurons from 2 rats, **Figure S 3**).

### Activation of LC^:PFC^ module is aversive and anxiogenic

Because activation of **LC^:PFC^** neurons did not change nociceptive thresholds we investigated whether activation of these neurons contributes to aversion or to the anxiety-like behaviour ascribed to the LC by McCall and colleagues (2015). We found that activation of **LC^:PFC^** caused avoidance of the drug-paired chamber in the CPP paradigm indicating a negative affective valence (**Figure 3**A).

**Figure 3:**
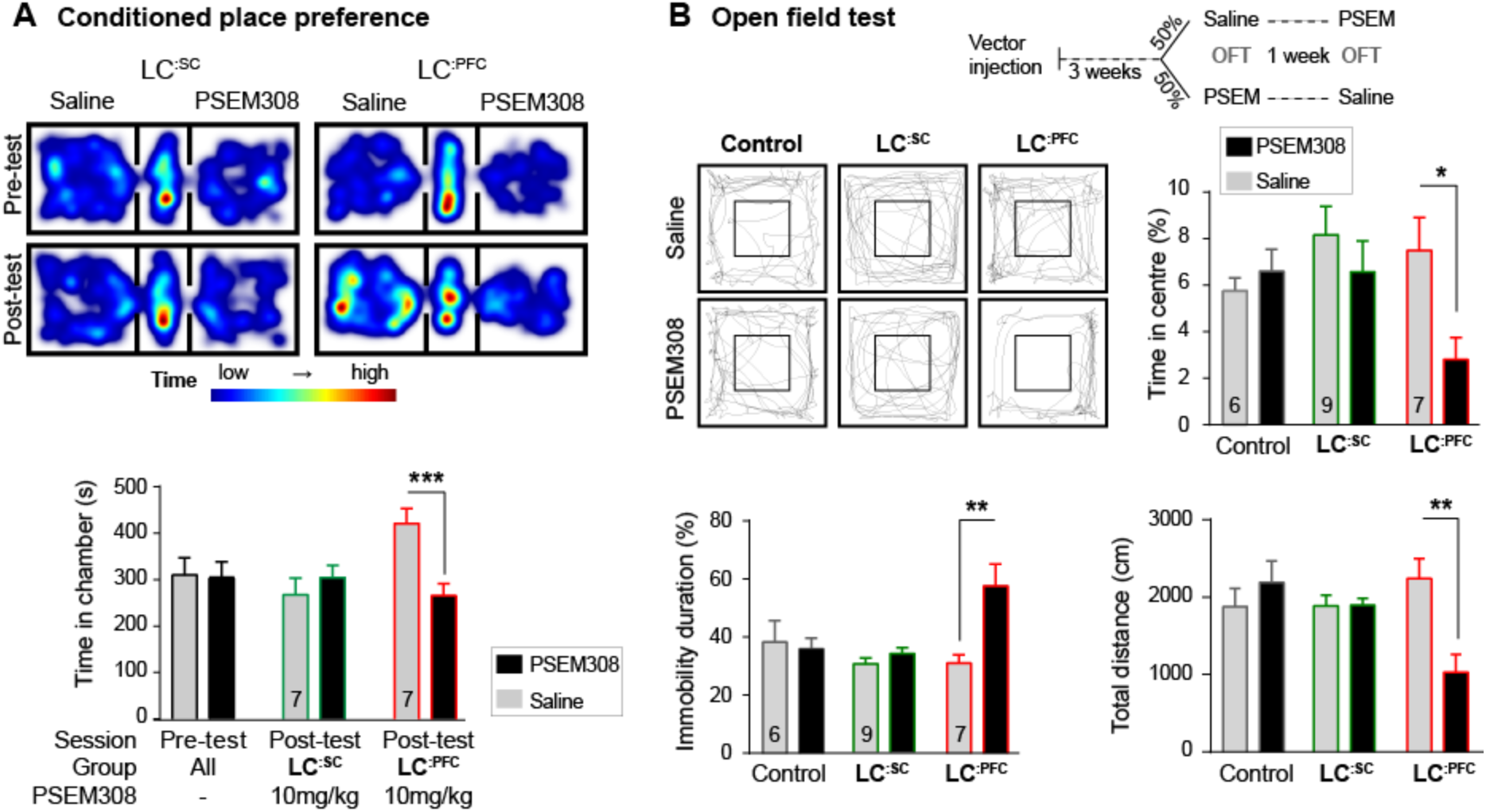
Chemogenetic activation of LC^:PFC^ neurons is aversive and anxiogenic unlike **LC^:SC^**. (A) Representative heat maps showing rat location in the place preference arena before and after conditioning for **LC^:SC^** (left) and **LC^:PFC^** (right) transduced rats (outer compartments 30x30cm). Activation of **LC^:PFC^** with PSEM308 (10 mg/kg) caused conditioned aversion to the paired chamber (2-way repeated measures ANOVA with Bonferroni’s multiple comparison between the conditioned and unconditioned chamber, *** P<0.001) and had no effect on **LC^:SC^** transduced rats. Pre-test data pooled for graphical presentation (B) Cross-over design for open field test (OFT) and tracks showing activity in the open field test (OFT) after PSEM308 (10 mg/kg) or saline (inner square 30x30 cm). PSEM308 reduced the time **LC^:PFC^** rats spent in the centre of the arena and increased the time these rats were immobile. In contrast, PSEM308 had no effect on **LC^:SC^** transduced rats (2-way RM-ANOVA with Bonferroni’s multiple comparison test, PSEM308 vs saline, * P<0.05, ** P<0.01).(See also Figure S 5)

The open field test uses the innate behaviour of rodents to avoid exposed areas as a measure of anxiety. Anxiogenic stressors typically reduce the time rodents spend in the centre of the arena and increase the time animals are immobile. Chemogenetic activation of **LC^:PFC^** reduced centre time by 62.6% and doubled immobility time in the open field test after PSEM308 i.p (**Figure 3**B). These findings indicate that chemogenetic activation of the **LC^:PFC^** neurons changes behaviour to produce anxiety/aversion and promoted immobility (as previously reported for high frequency stimulation of the whole LC (Carter et al., 2010)).

In contrast, when the **LC^:SC^** neurons were transduced, conditioning with PSEM308 was not aversive in the CPP paradigm (**Figure 3**A) and did not produce anxiety-like behaviour in the open field test (**Figure 3**B). There was no difference in the total distance animals travelled between drug or saline injection in the open field test for **LC^:SC^** animals (1894±90 PSEM308 vs 1885±137 cm saline, paired t-test p=0.94) suggesting there was no gross locomotor deficit. This indicates that chemogenetic activation of the **LC^:SC^** module does not produce the same negative affective behaviour that is seen with the **LC^:PFC^** module.

### Engagement of LC^:SC^ module attenuates neuropathic sensitisation

To test for possible analgesic benefit of chemogenetic activation of the **LC^:SC^** module in neuropathic pain, we employed the tibial nerve transection model (**Figure 4A** and **Figure S 4**). This procedure causes the progressive development of punctate mechanical (von Frey threshold lowered from 15.72±1.3 to 3.2±1.4 g, N=17) and cold hypersensitivity (increased frequency of paw withdrawals from 14.1±3.7 to 76.5±6.2 %, N=17) over 3 weeks (**Figure S 4**). Chemogenetic activation of the **LC^:SC^** module increased the mechanical withdrawal threshold after nerve injury (from 1.2±0.3 baseline to 11.7±3.3 g, t=30 min after PSEM308 i.p, N=6 **(Figure 4B**)). Similarly, we found activation of **LC^:SC^** produced a significant improvement in weight-bearing on the nerve-injured limb; linking the reduced evoked mechanical hypersensitivity to a spontaneous behaviour (**Figure S 4**). The chemogenetic analgesic effect on mechanical and cold allodynia was dose-dependent (1-10 mg/kg) and completely blocked by prior intrathecal (i.t) injection of the α2-adrenoceptor antagonist yohimbine (60 ng, 10.0±2.6 g PSEM308+vehicle to 1.2±0.4 g PSEM308+yohimbine, N=5 (**Figure 4D**)).

**Figure 4.**
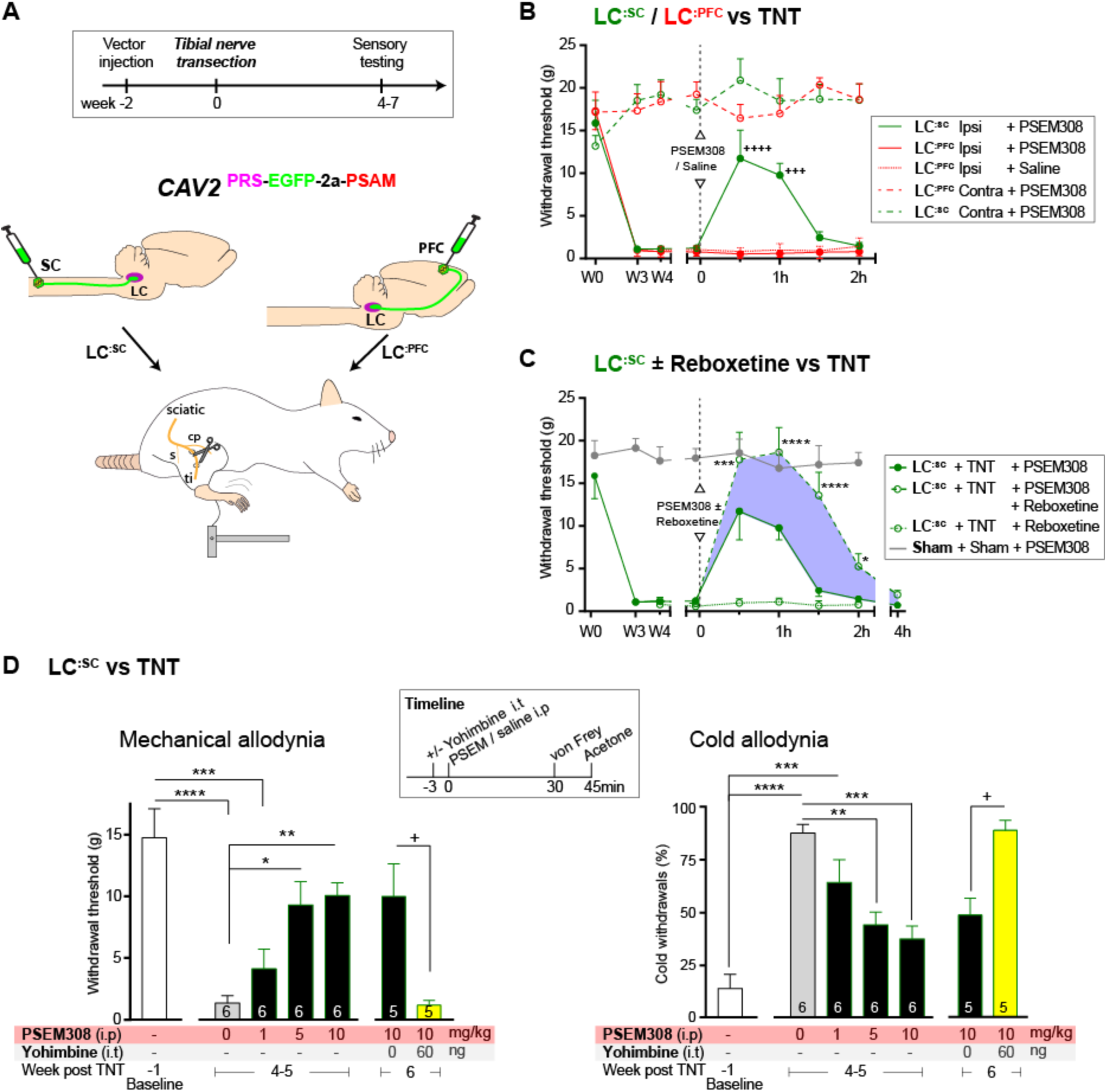
Chemogenetic activation of **LC^:SC^** is analgesic via a spinal α2-mechanism. (A) Experimental timeline of vector injection and the tibial nerve transection model of neuropathic pain (s = sural, cp = common peroneal and ti = tibial nerves). (B-C) Quantification of mechanical withdrawal threshold (von Frey, after Chaplan (1994)). (B) Nerve injured animals showed a mechanical sensitisation of the ipsilateral limb. Activation of **LC^:SC^** neurons (PSEM308 (10 mg/kg)) attenuated the neuropathic sensitisation (comparison vs baseline +++ P<0.001, ++++ P<0.0001, N=6 all groups). In contrast, activation of the **LC^:PFC^** had no effect on neuropathic sensitisation at any time point (C). Co-application of reboxetine i.t (1 mg/kg) enhanced the analgesic effect of **LC^:SC^** activation shown in panel (B). (D) Timeline of sensory testing in week 4-6. The analgesic effect of **LC^:SC^** activation on mechanical and cold allodynia is dose-dependent and is completely blocked by intrathecal yohimbine the α2-adrenoceptor antagonist (paired t-test, + P<0.05). All analysis by repeated measures ANOVA (1- or 2-way) with Bonferroni’s multiple comparison, * P<0.05, ** P<0.01, *** P<0.001, **** P<0.0001 unless otherwise stated (See also Figure S 4)

These findings indicate, that activation of **LC^:SC^** neurons is analgesic in this model of neuropathic pain and that the effect is mediated by noradrenaline acting on α2-adrenoceptors in the spinal cord. Therefore, we predicted that the noradrenaline reuptake inhibitor reboxetine (1mg/kg i.p), a dose too low to cause an analgesic effect on its own, would enhance chemogenetically-evoked **LC^:SC^** analgesia. Co-administration of reboxetine amplified and prolonged the analgesic effect of PSEM308 and completely reversed mechanical hypersensitivity (17.8±3.2 g) as compared to sham animals (17.9± 1.1g, (**Figure 4C**)).

In marked contrast to **LC^:SC^**, activation of the **LC^:PFC^** module by PSEM308 had either no effect or tended to increase evoked mechanical or cold responses in the nerve injury model and did not improve weight-bearing (**Figure 4**B and **Figure S 4**).

### Bidirectional modulation of spontaneous pain and pain-related behaviours by LC modules

We assayed the frequency of spontaneous foot-lifts as an index of ongoing pain caused by nerve injury. As anticipated, we found that nerve injury significantly increased the number of spontaneous foot-lifts over 5 min (6.2±0.7 sham, N=6 vs 30.7±3.9 TNT, N=17 (**Figure 5**B + **Video S 1**)). Activation of **LC^:SC^** more than halved the number of spontaneous foot-lifts in nerve injured rats (13.1±2.9), consistent with a reduction of ongoing pain. By contrast, activation of **LC^:PFC^** neurons almost doubled the number of foot-lifts (52.6±9.3 (**Figure 5**B)) suggesting a diametric modulation of spontaneous pain by these distinct subsets of LC neurons.

**Figure 5:**
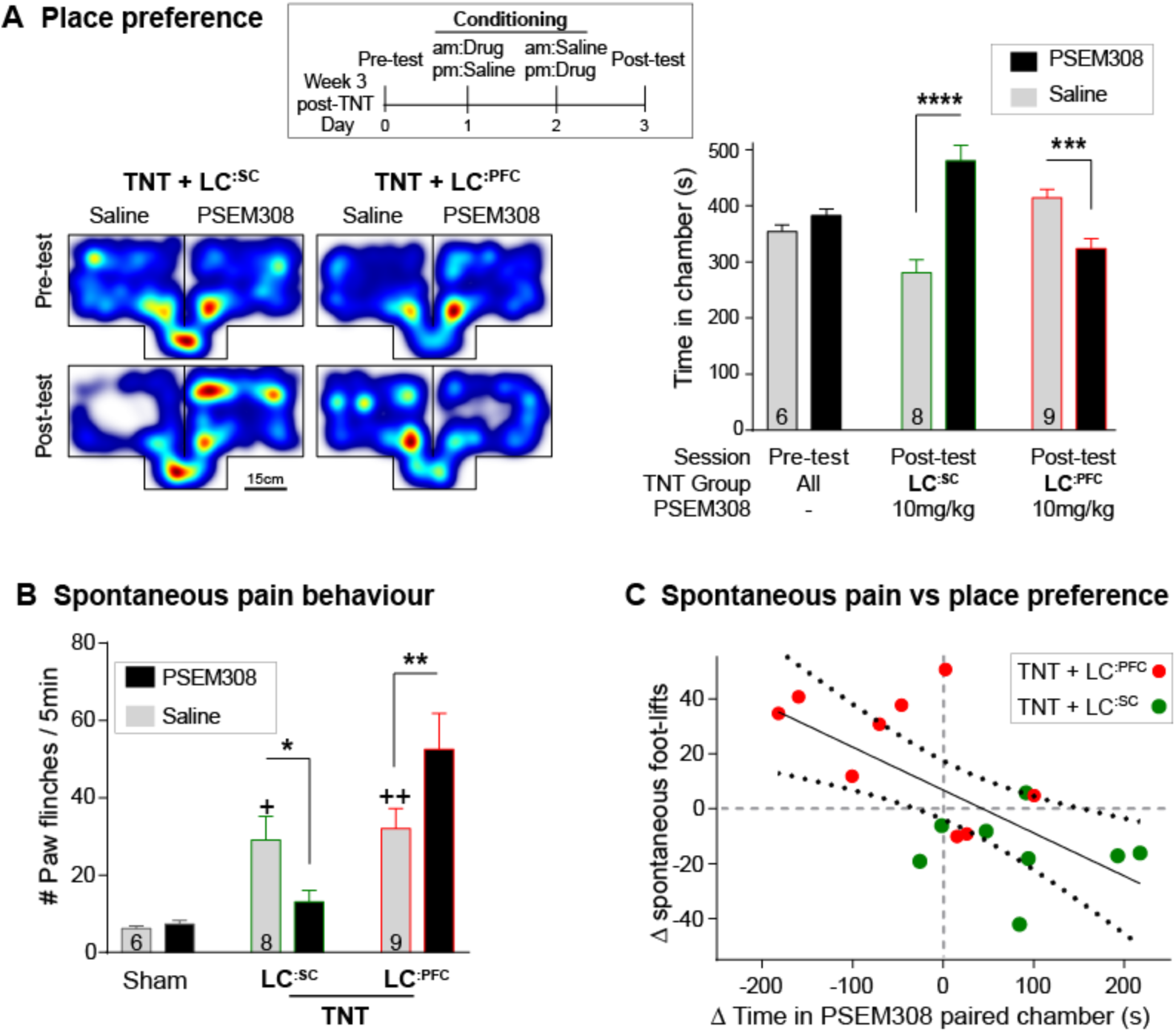
Chemogenetic activation of LC^:SC^ is rewarding in neuropathic pain model whilst activation of LC^:PFC^ neurons exacerbates spontaneous pain behaviour. (A) Timeline and representative location heat maps for a rat in CPP arena. In TNT animals activation of **LC^:SC^** now induces a robust place preference while activation of **LC^:PFC^** continues to produce aversion (2-way RM-ANOVA with Bonferroni’s multiple comparison between the conditioned and unconditioned chamber, ***P<0.001, ****P<0.0001). (B) TNT animals showed an increased frequency of paw flinches - a measure of spontaneous pain (saline treatment, RM-ANOVA with Bonferroni’s multiple comparison vs sham, + P<0.05, ++ P<0.01). Activation of **LC^:SC^** reduced and **LC^:PFC^** increased the frequency of flinches (2-way RM-ANOVA with Bonferroni’s multiple comparison PSEM308 (10 mg/kg) vs saline, * P<0.05, ** P<0.01). (C) Across both **LC^:SC^** and **LC^:PFC^** groups there was a correlation between the degree of modulation of spontaneous pain and the chamber preference (r= -0.65, P < 0.01). (See also Video S 1)

If this dichotomous action of the LC on spontaneous pain is also reflected in the affective experience of pain, then we anticipated that chemogenetic stimulation of the **LC^:SC^** module would reduce the negative pain-associated affect whilst **LC^:PFC^** activation should increase pain affect. In rats with neuropathic sensitisation, activation of **LC^:SC^** increased the time spent in the PSAM-paired chamber whilst **LC^:PFC^** activation induced aversion (**Figure 5**A). These data indicate that the neuropathic animals obtain benefit from **LC^:SC^** activation but that their situation is worsened by **LC^:PFC^** activation.

A link between the signs of ongoing pain and the affective component of pain is provided by the correlation between the PSEM308-induced modulation of chamber preference and the incidence of spontaneous foot-lifts across all groups of animals (**Figure 5**C). No correlation was found between chamber preference and PSEM308-induced changes in evoked responses (data not shown) suggesting that modulation of ongoing pain is a better predictor of the perceived benefit of this intervention.

The negative affect induced by **LC^:PFC^** activation was also evident from measures of anxiety in the open field test. Activation of **LC^:PFC^** reduced the time animals spent in the centre of the arena by 63% (5.4±1.6% saline to 2.0±0.7% PSEM308, N=7) and increased immobility by 139% compared to saline (47.3±6.0% saline to 65.9±7.7% PSEM308, N=7 (**Figure S 5**)). Furthermore, the effect on immobility caused by activation **LC^:PFC^** neurons was further enhanced by 135% by co-administration reboxetine (1 mg/kg) (61.3±8.0% PSEM308 to 82.8±3.9% PSEM308+reboxetine, N=6) indicating that the effect was caused by release of noradrenaline. In contrast, PSEM308 application had no effect on open field performance in **LC^:SC^** transduced or sham animals (**Figure S 5**).

## Discussion

By modifying LC cells using an intersectional vector strategy, based on transcriptional profile and projection territory, we rendered defined subgroups of noradrenergic neurons chemogenetically-activatable both *in vitro* and *in vivo*. Selective activation of the **LC^:SC^** or **LC^:PFC^** modules produced distinct changes in behaviour that were functionally antithetical in the context of pain states. The activation of the **LC^:SC^** neurons evokes an α2-adrenoceptor-mediated analgesic action that is sustained in a model of chronic neuropathic pain. This amelioration of the pain phenotype by **LC^:SC^** activation is seen for both evoked and spontaneous behaviour and was accompanied by reduced negative pain affect. In contrast activation of the **LC^:PFC^** produces an anxiety-like pattern of behaviour with negative affective valency and does not alter nociceptive sensitivity but instead increases expression of the signs of spontaneous pain after nerve injury. These data indicate that there is a modular functional organisation for these specific output targets of the LC and raises the question of whether this organisational principle is generalizable to other targets of the LC. Additionally, this approach in modulating a specific LC module establishes a proof of concept template for chronic pain treatment approaches with the potential for reduced side effects.

Within the pain field there has been a long interest in descending noradrenergic control which has been well characterised pharmacologically *in vitro* and *in vivo* (Millan, 2002, Yoshimura and Furue, 2006, Pertovaara, 2006, Jones, 1991). However, it has been difficult to selectively engage this system either clinically or in animal models because of its anatomical location, nuclear organisation, and lack of pharmacological targets to enable differentiated control with drugs. Here we use chemogenetic engineering approaches (Magnus et al., 2011) to manipulate and investigate the system with systemic drug administration. In targeting the **LC^:SC^** neurons we see an anatomical organisation that is similar to that previously described (Bruinstroop et al., 2012, Howorth et al., 2009a, Li et al., 2016, Westlund et al., 1983) with labelling in a cluster of around 15% of the LC neurons located in the ventral pole of the nucleus. **LC^:SC^** activation *in vivo* was anti-nociceptive in a heat withdrawal assay that was lateralised to the side of CAV2^PRS-EGFP-2A-PSAM^ vector injection in the spinal cord. Given that LC neurons were transduced bilaterally in the pons after this injection (as expected (Howorth et al., 2009a)), this indicates that there is a novel lateralised functional organisation for the control of nociception.

In a neuropathic pain model, activation of the **LC^:SC^** attenuated the degree of allodynia and hyperalgesia. This was seen for all sensory modalities tested and also applied to spontaneous measures of sensitisation i.e. foot-lifts. This benefit from engaging the **LC^:SC^** extends the findings from previous chronic intrathecal dosing studies with noradrenaline re-uptake inhibitors that also suppressed the neuropathic phenotype (Hughes et al., 2015). Given that we had documented a depletion of LC fibres in the spinal cord at the level of the nerve afferent input (Hughes et al., 2013) then it was not known whether there would be the potential to restore noradrenergic inhibition with the retrograde chemogenetic strategy. The demonstration of preserved actions indicates that both release of noradrenaline and the α2-adrenoceptor mechanism required are still functional. This is an important consideration for this kind of intervention as a therapeutic strategy; although we note that this strategy would have be successfully implemented after nerve injury (rather than transduced before nerve injury as here) if it were to be deployable clinically.

Pain is an emotive experience and relief of pain is a strong learning signal with positive affective valence i.e. animals display a preference for conditions where their pain is alleviated (King et al., 2009). Therefore, the preference of nerve-injured animals for with **LC^:SC^** activation indicates that this was analgesic rather than simply suppressing a reflex motor withdrawal – this parallels previous studies with intrathecal clonidine/noradrenaline re-uptake inhibitors that have shown a conditioned place preference (Hughes et al., 2015, Wei et al., 2013) or drive for self-administration (Martin et al., 2006). Additionally, no preference for **LC^:SC^** activation was seen in naïve rats, indicating that it is the analgesic action in the neuropathic model that has a positive valence rather than intrinsic rewarding effect of **LC^:SC^** stimulation. This is consistent with the sparse forebrain innervation (particularly of reward-related sites) for these **LC^:SC^** neurons (Howorth et al., 2009a, Li et al., 2016).

Engagement of the forebrain projecting **LC^:PFC^** produced a contrasting profile without any action on evoked-nociception and evidence of a facilitation of spontaneous pain behaviours in the neuropathic pain model. Additionally, activation of the **LC^:PFC^** was aversive and anxiogenic in both naïve and neuropathic animals. This aversive action agrees with previous studies of the LC (McCall et al., 2015) and may reflect high tonic levels of discharge produced by the chemogenetic activation approach. Such aversion and anxiety is a common component of chronic pain syndromes which likely contributes to the hypervigilance and catastrophizing. An increase in noradrenaline in the PFC has been shown in neuropathic pain models (Suto et al., 2014) that has been associated with an impairment of attention. Similarly, lesion studies of the **LC^:PFC^** have shown a reduction in pain associated anxiety and negative affect without any change in nociception (Bravo et al., 2016) which supports our finding that activation of **LC^:PFC^** produces negative affect and identifies this projection from the LC as a key player in the negative cognitive consequences of chronic pain.

These two opposing actions of the **LC^:SC^** and **LC^:PFC^** neurons to either suppress nociceptive input or to produce negative affective responses reflects the fact that they are distinct functional modules. This mirrors their anatomical segregation (Li et al., 2016) and implies that the modules could be capable of independent action, with recruitment in different behavioural contexts. We speculate that it may be the balance between recruitment of the *‘restorative’* **LC^:SC^** limb versus the *‘alarmist’* **LC^:PFC^** projection that determines whether pain following an injury resolves or persists into chronic pain (see also (Arora et al., 2016, Brightwell and Taylor, 2009, Taylor and Westlund, 2017)). Further the recruitment of either module may actually reciprocally inhibit the other given the α2-adrenoceptor-mediated feedback inhibition within the locus coeruleus (Aghajanian et al., 1977). This would also require each module to have different afferent drives that were independently regulated. At present, there is little evidence for such an organisation but this is an emerging territory for which the necessary neurobiological toolboxes are only now being developed (Schwarz et al., 2015). Their TRIO approach using CAV in combination with rabies vectors suggested that there was a distinction in the afferent innervation of the *‘medullary projecting’* LC neurons that we think likely represents collaterals from the **LC^:SC^**. This provides preliminary evidence that such an input–output specialisation may be present within the LC.

Previous studies have employed subtractive methods to study the role of the LC in pain regulation in animal models. These methods using toxins, knock outs or lesions to inhibit the noradrenergic system typically show subtle and sometimes contradictory effects on nociception (Martin et al., 1999, Jasmin et al., 2003, Jasmin et al., 2002, Tsuruoka and Willis, 1996, Howorth et al., 2009b). This may be in part ascribed to the long-term adaptation to the loss of a population of neurons. However, our findings of antithetical LC actions reported herein (and Hickey (2014)) provide an alternative explanation; namely that the LC is not a single functional entity with respect to pain regulation. Rather there are counterbalancing actions of individual modules and consequently the net effect of subtractive interventions will be difficult to predict/interpret. This consideration also applies to attempts to obtain therapeutic benefit from systemic dosing of noradrenaline re-uptake inhibitors in patients – while some benefit, a majority either obtain no benefit or suffer side effects that are often related to anxiety, changes in affect and sleep disturbances that are likely due to actions on ascending LC projections (Finnerup et al., 2015). Our findings indicate that targeting just the spinal projection could separate analgesia from many of the side effects.

In demonstrating this functional and anatomical dichotomy within the LC modules we have provided behavioural evidence that indicates that the LC has levels of sub-specialisation according to modalities. This builds upon prior work indicating anatomical and cellular specialisation within the LC (Loughlin et al., 1986, Kebschull et al., 2016, Chandler et al., 2014, Howorth et al., 2009a, Li et al., 2016). Given that the LC has been implicated in diverse neuronal behaviours such as learning and memory (Takeuchi et al., 2016, Martins and Froemke, 2015), strategy (Tervo et al., 2014), arousal (Carter et al., 2010) and anxiety (McCall et al., 2015). In each example the role of the LC in that behaviour has been considered as acting on a specific CNS substrate (i.e. hippocampus, auditory cortex, prefrontal cortex and amygdala) by inference somewhat in isolation. We speculate that the organisation of the LC outputs into modules allows these influences in different behavioural contexts to be independently selected and regulated. Given this organisational structure it may be that that the *Locus Coeruleus* would be more appropriately named *Loci Coeruleus* reflecting the fact that it appears to be a collection of noradrenergic modules each with distinctive paths and roles in regulating brain function.

## Materials and Methods

### EXPERIMENTAL MODEL AND SUBJECT DETAILS

All procedures conformed to the UK Animals (Scientific Procedures) Act 1986 and were approved by the University of Bristol local Ethical Review Panel. Animals were housed, with an enriched environment, under a reversed 12 h light/dark cycle at (23°C), with ad libitum access to food and water. All behavioural experiments were performed under red light conditions during the active dark phase of the cycle. Male Wistar rats from the University of Bristol colony were used for slice electrophysiology with vector injection at postnatal day 21 and subsequent use in experiments 10-14 days later. For behavioural assays and in vivo electrophysiology, male Wistar rats (120-200g, Charles River), randomly assigned to experimental groups, and injected with the appropriate vector. The observer was blind to both the experimental groups and injected solution for all behavioural experiments. All surgical procedures were carried out under aseptic conditions and animals were maintained at 37°C on a homeothermic pad. All animals in nerve injury experiments were single housed.

### METHOD DETAILS

#### Generation of LV^PRS-EGFP-2A-PSAM^

The chemogenetic excitatory ionophore PSAML141F,Y115F:5HT3HC (PSAM) (Magnus et al., 2011) was obtained in inverted orientation from Addgene (#32477). This sequence information was used to 1) generate a CMV promotor driven expression plasmid by excising the PSAM-IRES-EGFP cassette with BsrGI/NheI and subsequent ligation into the pre-cut expression plasmid (BsrGI/NheI) which caused the flip in sense orientation and 2) to design (using ApE) a HA tagged (8 C-terminal amino acids) version of PSAM in the EGFP-2A-PSAM_HA_ expression cassette with the objective of producing stronger EGFP fluorescence whilst PSAM receptor function is no different to the original cassette (**Figure S 1**). The CMV-EGFP-2A-PSAM_HA_ cassette was custom synthesised in a plasmid (pCMV-EGFP-2A-PSAM_HA_) by GeneArt AG (Sequence available on request).

The LV was produced as previously detailed (Coleman et al., 2003, Hewinson et al., 2013). In short, the LV is based on a HIV-1 vector pseudotyped with vesicular stomatitis virus glycoprotein. The transgene product EGFP-2A-PSAM_HA_ was excised with BamHI and BstBI and ligated into a pre-cut lentiviral (pTYF) backbone vector harbouring the noradrenergic specific PRSx8 (PRS) promotor (Hwang et al., 2001) to generate pTYF-PRS-EGFP-2A-PSAM_HA_-WPRE. The LVs were produced by cotransfection of Lenti-X293T cells (Clontech) with the shuttle vector pTYF-PRS-EGFP-2A-PSAM_HA_-WPRE or pTYF-PRS-EGFP-WPRE, along with a packaging vector, pNHP, and the envelope plasmid pHEF-VSVG. The virus containing supernatant was collected on three consecutive days post transfection, pooled and concentrated by ultracentrifugation. The titre, measured in transducing units (TU/ml), was assayed by establishing the infection rate of an LV produced in parallel and harbouring the expression cassette for placental alkaline phosphatase (Hewinson et al., 2013). The LV was used for microinjection at a titre of 7.5x10^10^ TU/ml for LV-PRS-EGFP-2A-PSAM_HA_ and at 3.7x10^10^ TU/ml for LVPRS-EGFP.

#### Generation of CAV2^PRS-EGFP-2A-PSAM^

The EGFP-2A-PSAM_HA_ cassette was excised from the custom synthesised plasmid (pCMV-EGFP-2APSAM_HA_) with ApaI and HpaI. The CAV2 shuttle vector pTCAV2-PRS-ChR2-mCherry (previously described (Li et al., 2016)) was cut with ApaI/HpaI to remove ChR2-mCherry and subsequently ligated to generate pTCAV2-PRS-EGFP-2A-PSAM_HA_. The resulting plasmids were then purified and digested with BamHI/NotI for homologous recombination into the SwaI linearized CAV2 genome (pCAVΔE3Sce) in BJ5183 cells (Agilent Technologies) following the manufacturer’s protocol. CAV2 vectors were produced using previously described methods (Ibanes and Kremer, 2013). Briefly, DKSceI dog kidney cells constitutively expressing the fusion protein of the endonuclease SceI and the estrogen receptor (SceI-ER) were grown in complete DMEM (with 10% fetal bovine serum, nonessential amino acids, L-glutamine, penicillin and streptamycin and G418) on 10cm dishes to 80% confluency. Subsequently, cells were treated with 1mM OH-tamoxifen for 4h to induce translocation of SceI-ER into the nucleus. The tamoxifen pre-treated DK-SceI cells (8x10^6^) were seeded in 10cm dishes containing 1mM OH-tamoxifen and immediately transfected with 15 μg of the supercoiled pCAV genome using 30μl turbofect. The transfection medium was replaced by complete DMEM the following morning. Cells were harvested and virus released by freeze-thaw cycles to prepare initial virus containing stocks for vector amplification in DK-SceI cells (Ibanes and Kremer, 2013). Vector stocks were titrated in functional transducing units (TU) per ml as previously described (Li et al., 2016, Ibanes and Kremer, 2013, Kremer et al., 2000) and diluted to 0.6x10^12^ TU before injection.

#### Stereotaxic injections

The procedures for viral vector injections into the LC, the PFC or the SC have been described in detail previously (Hickey et al., 2014, Howorth et al., 2009a, Li et al., 2016). In brief, rats were anesthetized by intraperitoneal injection of ketamine (5 mg/100 g, Vetalar; Pharmacia) and medetomidine (30 μg/100 g, Domitor; Pfizer) until loss of paw withdrawal reflex. The rat was placed in a stereotaxic frame. Burr holes and microinjections were made using a Drill and Injection Robot with Wireless Capillary Nanoinjector (NEUROSTAR, Germany).

Locus coeruleus targeting in rat pups (p19-21): A dorso-ventral injection track was started at 1mm lateral and 1mm posterior to lambda with 10° rostral angulation. Four injections of 0.25μl of vector were made at four locations beneath cerebellar surface (-4.6mm, -4.9mm, -5.2mm and -5.5mm).

Locus coeruleus targeting in adult rats (120-200g): A dorso-ventral injection track was started at 1.2mm lateral and -2.1mm posterior to lambda with 10° rostral angulation. Three injections of 0.3μl were made at -5.3mm, -5.5mm and 5.8mm depth. In rats intended for in vivo LC recordings a screw was inserted into the burr hole to simplify later CNS access.

Prefrontal cortex: Four burr holes were drilled bilaterally, 0.7mm lateral to the midline, at +2mm, +1.2 mm, +0.8 mm and -0.2mm from bregma. At each site 0.2μl of vector was injected at 1.0mm and 1.5mm depth to the brain surface. At +2mm further injections were made at -5 and -5.5 mm depth from the brain surface

Spinal cord: The vertebral column was exposed and clamped at vertebral segment L3. A laminectomy was performed at T12-L1 and 0.4μl CAV2 vector injected bilaterally into the L3-L4 segment of the spinal cord (SC) at six injection sites in total. These injections were located 0.5mm lateral from the midline and 0.5mm deep and spread in pairs over the segmental extent. *For unilateral injections (pain studies)* six injections of (0.4μl) were made unilaterally into the right lumbar dorsal horn.

#### Electrophysiology

##### Patch clamp recordings from acute pontine slices

Pontine slices were prepared from juvenile (age p28-p32) Wistar rats 7–14 days after lentiviral transduction as previously described (Hickey et al., 2014). Rats were deeply anesthetized with isoflurane 5%, decapitated and the brain quickly removed and immediately chilled in ice-cold cutting solution (similar to the recording solution but NaCl was reduced from 126mM to 85 mM and substituted with sucrose 58.4mM) and the brainstem blocked. Horizontal slices (300–350μm thick) of the pons were cut from dorsal to ventral using a vibratome (Linearslicer Pro 7; DSK) in cold (4°C) cutting solution. After cutting, slices were kept at room temperature in carbogenated recording solution (NaCl 126 mM, KCl 2.5 mM, NaHCO_3_ 26 mM, NaH_2_PO_4_ 1.25 mM, MgCl_2_ 2 mM, CaCl_2_ 2 mM, and D-glucose 10 mM saturated with 95%O_2_/5%CO_2_, pH7.3, osmolality 290mOsm/L) for at least 1 h to recover before recording. Pontine slices were transferred into the recording chamber of an upright fluorescence microscope (DMLFSA; Leica Microsystems), superfused with artificial CSF at a rate of 4–8 ml/min heated to 35°C. Borosilicate glass patch pipettes (Harvard Apparatus GC120F-10, resistances of 4–6MΩ) were filled with internal solution (K-gluconate 130 mM, KCl 10 mM, Na-HEPES 10 mM, MgATP 4 mM, EGTA 0.2 mM, and Na_2_GTP 0.3 mM). Cells were identified under gradient contrast illumination and examined for transduction by epifluorescence illumination. Whole cell voltage and current clamp recordings were obtained with a Multiclamp 700A amplifier (Axon instruments). All membrane potentials were corrected for a junction potential of 13 mV. Drugs were infused in the bath solution. Spike2 software (CED, Cambridge Electronic Design) with customized scripts was used to acquire and store data.

##### In vivo extracellular recordings from the locus coeruleus and local application of PSEM308

Anaesthesia was induced with 1.2-2 g/kg urethane (Sigma) i.p to achieve loss of paw withdrawal reflex. Once induced, urethane anaesthesia typically remained stable for 5-7 hours. The animal was placed in a stereotaxic frame, the skull exposed and the screw above the viral injection site removed to gain access to the CNS. In vivo single units were recorded with preloaded quadruple-barrelled microelectrodes (Kation Scientific, Carbostar-4 recording/iontophoresis microelectrode, E1041 standard) connected to a drug injector (Parker, Picospritzer II). Recordings were referenced against a silver chloride electrode pellet that was inserted between muscles and skin near the recording site. Neuronal activity was recorded via a Multiclamp 700A amplifier (with #CV-7b headstage, Axon instruments) filtered at 3 kHz and digitized at 10 kHz using a 1401micro and Spike2 software (CED, Cambridge Electronic Design) with custom scripts was used to store data and to control the drug delivery. Locus coeruleus cells were identified by their wide (~1 ms), large amplitude action potentials, the slow spontaneous firing frequency (0.5-5Hz) and a characteristic burst of action potentials in response to contralateral hind-paw pinch (Hickey et al., 2014, Cedarbaum and Aghajanian, 1976). Action potential discharge frequency was measured in 5 s bins and compared before, during, and after drug application. The average depth of electrode placement for LC recordings was 5.7±0.14 mm from the cerebellar surface.

##### In vivo single unit recordings from the dorsal horn

Animals were terminally anaesthetised with urethane (1.2-2 g/kg i.p, Sigma) The spinal cord was exposed by laminectomy over the L1-L4 spinal segment for recordings (Funai et al., 2014, Furue et al., 1999). The animal was placed in a stereotaxic frame and the spinal cord stabilised with two spinal clamps at T1 and L5 and a bath was formed by skin elevation. Warmed (36°C), Krebs solution was used to continually superfuse the spinal cord (NaCl 117 mM, KCl 3.6 mM, NaHCO_3_ 25 mM, NaH_2_PO_4_ 1.2 mM, MgCl_2_ 1.2 mM, CaCl_2_ 2.5 mM and D-glucose 11 mM saturated with 95%O_2_/5%CO_2_, pH7.3, osmolality 290mOsm/L). The dura mater was removed under binocular vision (Leica MZ6). A reference electrode (World Precision Instruments, Ag/AgCl Electrode pellets, EP2) was inserted into the muscle layer near the recording area and spinal units were recorded with a stainless-steel microelectrode (FHC, UESSEGSEFNNM, 8-8.8MΩ) that was positioned ~500 μm from the midline. Neuronal activity was amplified and low pass filtered at 3 kHz using a Multiclamp 700A amplifier (Axon instruments with #CV-7b headstage). The data were digitized at 10 kHz (CED Power1401) and stored using Spike2 software (CED, Cambridge Electronic Design) with customized scripts to monitor pinch force applied from custom calibrated forceps with tip pads of area 3.5mm^2^. These forceps were used to ensure that the pinch stimulus was standardised between applications and across animals. The forceps were equipped with strain gauges and a pinch pressure of 70 g mm^-2^ (Drake et al., 2014) was used throughout all experiments to compare pinch-evoked neuronal activity (NB this pressure was above the threshold for pain when applied to the experimenter’s index finger).

Wide dynamic range (WDR) neurons were identified in the deep dorsal horn by graded responses to both non-noxious touch and noxious pinch. Their discharge frequency was measured in response to light touch with a brush or to sustained pinch pressure over five second time bins immediately after stimulus application.

##### Tibial nerve transection model

Rats were anesthetized by intraperitoneal injection of ketamine (5 mg/100 g, Vetalar; Pharmacia) and medetomidine (30 μg/100 g, Domitor; Pfizer) until loss of paw withdrawal reflex. A 1.5-2cm long skin incision was made in the right hind leg in line with the femur (Pertin et al., 2012). The fascial plane between the gluteus superficialis and the biceps femoris was located and blunt dissected to expose the three branches of the sciatic nerve (i.e. common peroneal, sural and tibial nerves). The tibial nerve was carefully exposed. Subsequently, two tight ligations (5-0 silk) were made approximately 2mm apart and the neve segment in between transected and removed. The nerve and muscle were placed back into the correct position and the skin sutured. For sham surgery, the same procedure without ligation and transection of the tibial nerve was performed.

##### Intrathecal injection after nerve injury

The injection procedure was similar to a previous report (De la Calle and Paino, 2002) Rats, 6 weeks post TNT (400-500g), were briefly induced with isoflurane in medical oxygen before transferring into a face mask with 1.5-2% isoflurane. The rat was placed in prone position on a 5cm tall plastic board so that the hind legs hang over the board to arch the lower back and to open the intervertebral space between L4-L5 spinal segments. Subsequently, a 25G needle was inserted transcutaneously to the vertebral canal (typically producing a tail twitch) and 10-20ul of vehicle or yohimbine (3μg/μl in sterile saline with 20% DMSO) injected. PSEM308 was injected i.p immediately after recovery from anaesthesia and sensory testing began 30min thereafter.

As previously described (Hughes et al., 2013), we observed that the mechanical withdrawal thresholds fell in the contralateral paw after intrathecal injection of yohimbine in nerve injured animals. This phenomenon was used as an internal control for successful intrathecal targeting because it was more reliable than a tail flick response (**Figure S 4**).

##### Behaviour

Rats underwent two 30 minute habituation sessions (unless otherwise stated) on consecutive days before testing. Animals were placed into Plexiglas chambers 20 minutes before sensory assessment began on the test day. All experiments, apart from conditioned place preference (CPP), were conducted in a cross-over design comparing saline and PSEM308 (10 mg/kg, unless otherwise specified). The experimenter was always blind to the animal group and to the administered solution.

##### Hargreaves’ Test

Thermal withdrawal latencies were measured for the hind paw using the method described by Hargreaves et al. (1988). Animals were placed into a Plexiglas chamber on top of a glass plate. The radiant heat source (Ugo Basile Plantar test) was focused onto the plantar surface of the hind paw and the time to withdrawal recorded from 40 min before and up to 60 minutes after intraperitoneal injection of the selective agonist PSEM308 or saline. Each withdrawal value was the mean of 2 tests and tests were repeated every 15 minutes. A 24 second cut-off value was used to avoid tissue damage. The heat source intensity was adjusted so that control rats responded with a latency of 7-8 seconds.

##### Spontaneous foot lifts

Animals were placed into a Plexiglas chamber on a metal grid and a webcam (Logitech HD Pro Webcam C920) positioned underneath. The number of spontaneous footlifts and flinches was recorded over a 5 minute period 20 minutes after i.p injection of PSEM308 / saline.

##### von Frey test

To assess punctate mechanical sensitivity animals were placed into a Plexiglas chamber on a metal grid and von Frey filaments (0.4g-26g) were applied to the lateral aspect of the plantar surface of the hind paw. The 50% paw withdrawal threshold was determined using the Massey-Dixon up-down method (Chaplan et al., 1994). This procedure was repeated every 30 minutes starting 1h prior and up to 4h after i.p injection of PSEM308 (1, 5 and 10 mg/kg) or saline.

##### Acetone Test

To assess cold sensitivity animals were placed into a Plexiglas chamber on a metal grid and an acetone drop on top of a 1ml syringe was presented to the hind paw from underneath 5 times, 2 minutes apart. Data are expressed as the percentage of withdrawals. Animals were tested prior to and 45 minutes after i.p injection of PSEM308 or saline.

##### Incapacitance Test

The distribution of weight between the hind-limbs was measured with an incapacitance tester (Linton Instrumentation). Animals were habituated to the Plexiglas chamber with their hind paws on two force transducers in 4 sessions (each lasting 3 minutes) on two consecutive days. Measures were taken 45 minutes after injection of PSEM308 or saline. The percentage of weight borne on the injured leg was averaged over these 4 measures.

##### Conditioned place preference/aversion

All rats were habituated to an unbiased three-compartment box, which contained a central neutral chamber with different association chambers either side. The outside compartments have visual and tactile cues e.g. “bars” or perforated “holes” as flooring and differently patterned (but equal luminosity) wallpapers to maximise differentiation between the compartments. On the first day of the experiment animals were allowed to roam freely through all three compartments and a Basler camera (acA1300-60gm) with a varifocal lens (Computar H3Z4512CS-IR) connected to EthovisionXT (Noldus) was used to record the time rats spent in either of the outer compartments as a baseline value. Subsequently, rats underwent four conditioning sessions in which the outside compartments were paired with PSEM308 injection or saline injection. Two pairing sessions per day at least 4h apart on two consecutive days were conducted in a counterbalanced design. For each conditioning session the animals were placed into the locked outer compartment 5 minutes after the i.p injection and left for 30min. On the test day, the doors between the compartments were opened and the time spent in the outer compartments measured once more. It is expected that the choice of the animal’s preferred environment in the post-test is influenced according to the valence of the chemogenetic stimulation. Noldus EthoVision^®^ XT was used to analyse recorded videos and to quantify the time animals spent in the paired chambers.

##### Open field test

On the first day of testing animals were randomly assigned into two groups and received PSEM308 or saline 20-25 minutes before placing the subjects into the open field arena (56.5x56.5cm with an inner zone of 30x30cm) for 6 minutes. Exploration behaviour (distance travelled, location in central / perimeter areas and immobility (velocity < 1.75cm/s)) was tracked for 5 minutes starting 60 seconds after entering the arena using EthoVisionXT. One week later the open field experiment was repeated using the other solution.

#### Histology

##### Tissue Collection and Processing

Rats were killed with an overdose of pentobarbital (Euthatal, 20 mg/100 g, i.p; Merial Animal Health) and perfused transcardially with 0.9% NaCl (1 ml/g) followed by 4% formaldehyde (Sigma) in phosphate buffer (PB; pH 7.4, 1 ml/g). The brain and spinal cord were removed and post-fixed overnight before cryoprotection in 30% sucrose in phosphate buffer. Coronal tissue sections were cut at 40μm intervals using a freezing microtome and left free floating for fluorescence immunohistochemistry.

##### Immunofluorescence

Tissue sections were blocked for 45minutes in phosphate buffer containing 0.3% Triton X-100 (Sigma) and 5% normal goat serum (Sigma). Incubated on a shaking platform with primary antibodies to detect EGFP (1:3000), DBH (1:2000), or HA tag (1:500-1:1000) for 14–18 h at room temperature. After washing, sections were then incubated for 3 h with appropriate Alexa Fluor^®^ secondary antibodies. A Leica DMI6000 inverted epifluorescence microscope equipped with Leica DFC365FX monochrome digital camera and Leica LAS-X acquisition software was used for widefield microscopy. For confocal and tile scan confocal imaging, a Leica SP5-II confocal laser-scanning microscope with multi-position scanning stage (Märzhäuser, Germany) was utilized. Images were exported as TIF files before being processed, analysed and prepared for presentation using Fiji-ImageJ software.

Stereological cell counts (Miller et al., 2014) were made by stepping through a confocal z-stack with 1μm spacing and EGFP positive cells were only counted if the top of the soma was unambiguously located within the imaged dissector volume of the section (12μm). The average section thickness after histological processing was 18.0±0.7μm (30 sections from 3 animals) and dual guard zones of 2μM were applied. Therefore, the height sampling factor was 0.66 (hsf = dissector height/average section thickness). The section sampling factor was 0.25 (ssf = 1 / number of serial sections). The correction was made using the following formula.

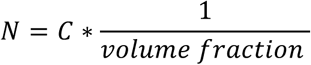

Where N is the estimated number of cells and C the sum of all counted cells. The volume fraction is the product of ssf, hsf and asf (area sampling fraction). Because all EGFP positive neurons in the LC area were counted the area sampling fraction was omitted from the volume fraction.

### QUANTIFICATION AND STATISTICAL ANALYSIS

All statistical analysis was conducted using GraphPad Prism 7. All data are presented as mean ± SEM (unless otherwise specified). Sample size estimates were based either on previous data from similar experiments (Hughes et al., 2015, King et al., 2009, McCall et al., 2015) or for novel experiments based on an initial pilot experiment to obtain an estimate of effect size and variance for a power calculation with alpha <0.05 and beta 0.8 (using G*Power) to inform subsequent definitive experiments. T-tests, one-way or two-way ANOVA were used to compare groups as appropriate. Bonferroni’s post-test was used for all comparisons between multiple groups. Differences were considered significant at p < 0.05. The number of replications (n) equals the number of cells for electrophysiological experiments (the number of individual rats is stated in the relevant section of the text). Whereas (N) is the number of animals tested in behavioural assays and the number of LCs that were analysed in histological experiments. Each experiment was repeated with at least two independent batches of animals for each condition.

Exclusions: One animal was excluded from conditioned place preference experiments because it learnt to escape from the arena. Data from four spinal cord WDR neurons were excluded because the recording was lost before the end of the protocol. One animal (in the **LC^:PFC^** group) was excluded from analysis because of a failure of vector transduction discovered on post hoc histology.

### RESOURCE TABLE

**Table S1:**
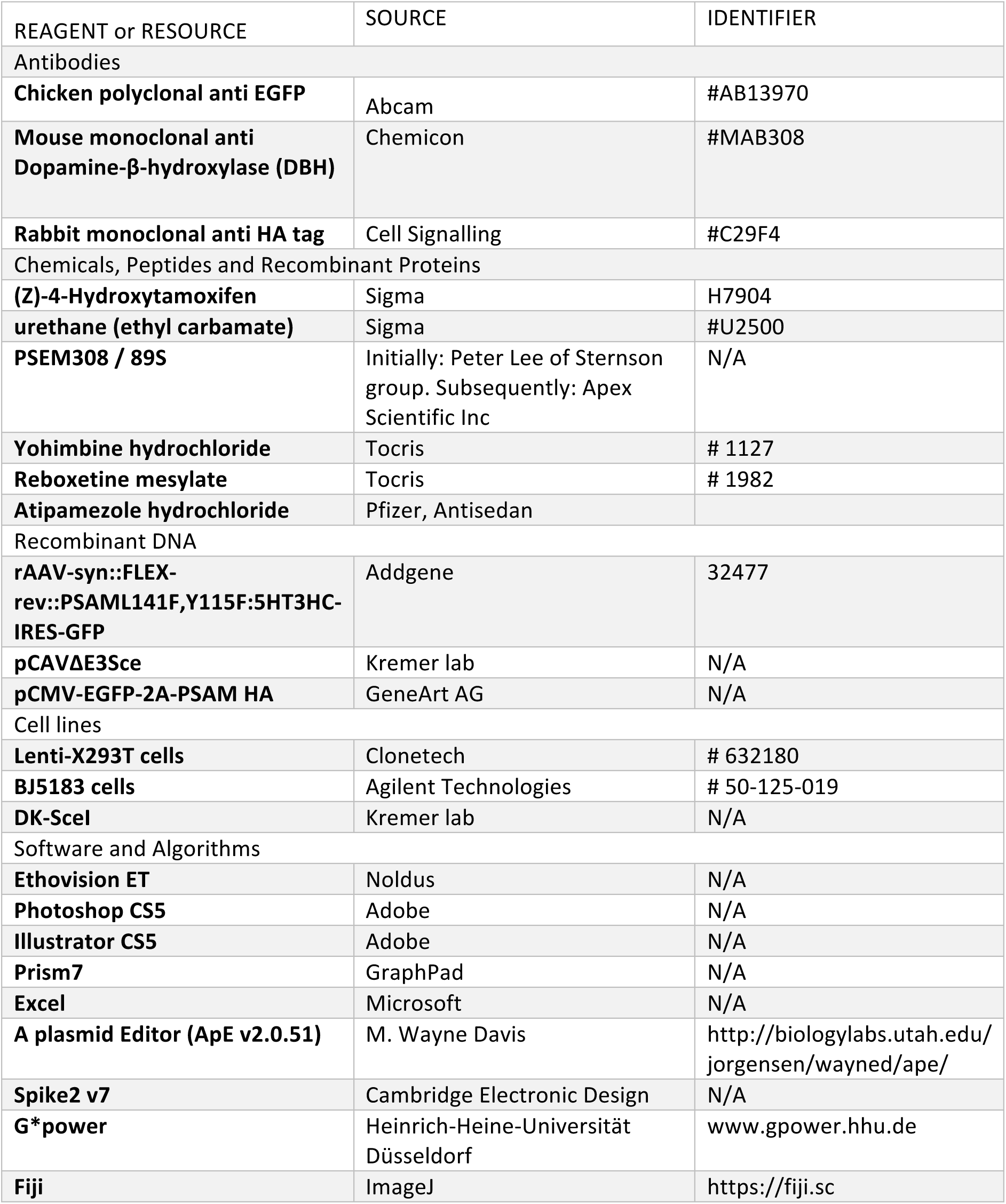
Key resource table

| REAGENT or RESOURCE | SOURCE | IDENTIFIER |
| --- | --- | --- |
| **Antibodies** |  |  |
| **Chicken polyclonal anti EGFP** | Abcam | #AB13970 |
| **Mouse monoclonal anti Dopamine-β-hydroxylase (DBH)** | Chemicon | #MAB308 |
| **Rabbit monoclonal anti HA tag** | Cell Signalling | #C29F4 |
| **Chemicals, Peptides and Recombinant Proteins** |  |  |
| **(Z)-4-Hydroxytamoxifen** | Sigma | H7904 |
| **urethane (ethyl carbamate)** | Sigma | #U2500 |
| **PSEM308 / 89S** | Initially: Peter Lee of Sternson group. Subsequently: Apex Scientific Inc | N/A |
| **Yohimbine hydrochloride** | Tocris | # 1127 |
| **Reboxetine mesylate** | Tocris | # 1982 |
| **Atipamezole hydrochloride** | Pfizer, Antisedan |  |
| **Recombinant DNA** |  |  |
| **rAAV-syn::FLEX-rev::PSAML141F,Y115F:5HT3HC-IRES-GFP** | Addgene | 32477 |
| **pCAVΔE3Sce** | Kremer lab | N/A |
| **pCMV-EGFP-2A-PSAM HA** | GeneArt AG | N/A |
| **Cell lines** |  |  |
| **Lenti-X293T cells** | Clontech | # 632180 |
| **BJ5183 cells** | Agilent Technologies | # 50-125-019 |
| **DK-Scel** | Kremer lab | N/A |
| **Software and Algorithms** |  |  |
| **Ethovision ET** | Noldus | N/A |
| **Photoshop CS5** | Adobe | N/A |
| **Illustrator CS5** | Adobe | N/A |
| **Prism7** | GraphPad | N/A |
| **Excel** | Microsoft | N/A |
| **A plasmid Editor (ApE v2.0.51)** | M. Wayne Davis | http://biologylabs.utah.edu/jorgensen/wayned/ape/ |
| **Spike2 v7** | Cambridge Electronic Design | N/A |
| **G\*power** | Heinrich-Heine-Universität Düsseldorf | www.gpower.hhu.de |
| **Fiji** | ImageJ | https://fiji.sc |

### CONTACT FOR REAGENT AND RESOURCE SHARING

Further information and requests may be directed to, and will be fulfilled by, the corresponding author and Lead Contact Dr. Anthony E. Pickering. Plasmid sequences are available on request

## Acknowledgements

This work was supported by:

- Wellcome Trust Senior Clinical Research Fellowship (gr088373) – A.E.P.
- University of Bristol Postgraduate Research Scholarship – S.H.
- EMBO short-term fellowship programme to facilitate researcher exchange (S.H.) between Kremer and Pickering lab

The authors thank:

- *The Sternson lab at Janelia Research Campus* for kind gifts of PSEM89s/PSEM308 for initial trials, plasmids containing PSAM sequences and for technical advice
- Hidemasa Furue and Robert Drake for advice on electrophysiological experiments and data presentation
- Sandy Ibanes for help with vector production

## ADDITIONAL RESOURCES

### Supplementary information

**Figure S 1:**
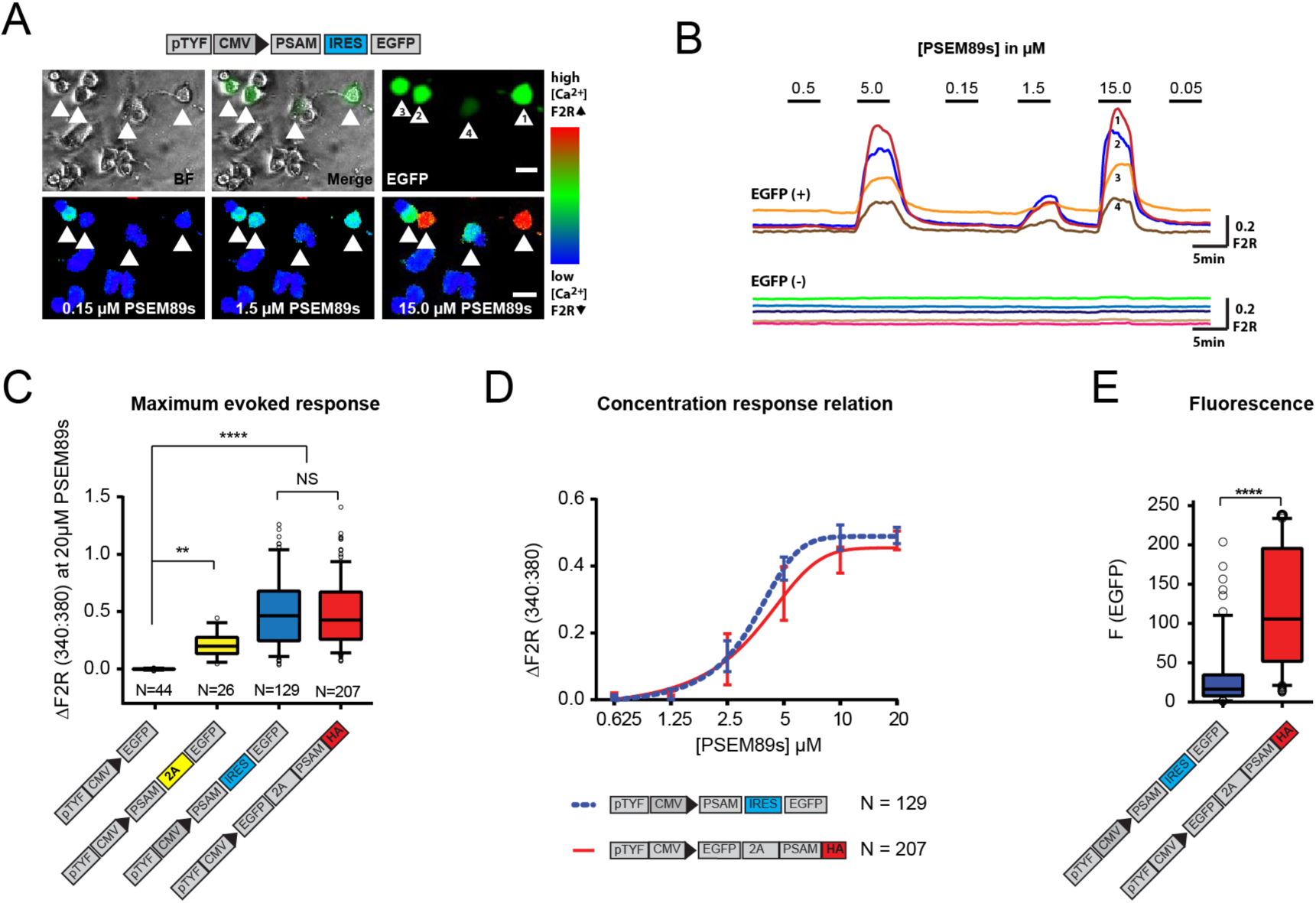
Related to Results: Targeting and activation of noradrenergic LC neurons with PSAM. Generation of a traceable version of the PSAM in an expression cassette with enhanced fluorescence (A) Schematic of the PSAM (PSAM L141F,Y115F:5HT_3_HC) and EGFP co-expression plasmid. Top row shows 4 EGFP+ transfected and ten EGFP-PC12 cells. The bottom row shows [Ca^2+^]_i_ increases in the transduced cells to the selective agonist PSEM89s. [Ca^2+^]_i_ measured by Fura2 340:380 fluorescence ratio - F2R. (B) [Ca^2+^]_i_ responses to PSEM89s in 4 EGFP+ and 5 EGFP-PC12 cells. (C) The maximum Ca^2+^ transient evoked by PSEM89s is sensitive to the length of C-terminal tag on PSAM. 18 amino acids remain at the C-terminal end of PSAM when the 2A peptide is downstream of PSAM. This C-terminal residue reduced the maximum PSEM89s-evoked response. There was no difference between the original PSAM-IRES-EGFP plasmid and the HA tagged expression cassette EGFP-2A-PSAMHA (p<0.0001 Kruskal Wallis test with Bonferroni’s multiple comparison NS p>0.05, **p<0.01, ****p<0.0001). (D) The C-terminal HA tag does not interfere with the concentration response relationship (p=0.6384 sumof-squares F test). EC50: PSAM-IRES-EGFP - 3.37±0.04 vs EGFP-2A-PSAM_HA_ - 3.86±0.09 μM (E) The upstream EGFP-P2A sequence produces a significantly brighter fluorescent signal than the IRES linker (Mann-Whitney test p<0.0001). Calcium transients are shown as mean ±SEM and fluorescence data as median with interquartile intervals. N equals the number of cells analysed from at least three replicated experiments.

**Table S2: Related to Figure 1.** LC neuronal electrophysiological properties obtained from whole cell patch clamp recordings are not significantly different between transduced and non-transduced cells. In current clamp, spontaneous discharge frequency was calculated over 1 minute. The threshold for action potential initiation was defined as the point where velocity of depolarisation exceeded 7.5Vs^-1.^ Threshold was used as the reference to measure spike and afterhyperpolarization amplitude. Action potential duration was measured at half spike amplitude. The duration of after-hyperpolarization was defined as the trough-width below resting potential. Data from 25 action potentials were averaged per cell. Input resistance (R) was measured in voltage clamp from the average of 10 current responses (I) to a 10mV membrane hyperpolarization (V). Data presented as mean±SEM, and P values from unpaired t-test.

|  | Non-transduced LC neurons | Transduced LC neurons | P value |
| --- | --- | --- | --- |
| n= | 7 | 7 |  |
| Resting Potential (mV) | -59.9±2.0 | -60.4±1.8 | 0.85 |
| Firing rate (Hz) | 1.85±0.45 | 1.12±0.43 | 0.27 |
| Threshold (mV) | -47.0±2.9 | -48.4±1.5 | 0.70 |
| AP Amplitude (mV) | 59.7±2.3 | 62.4±2.6 | 0.44 |
| AP Duration (ms) | 1.63±0.25 | 1.43±0.08 | 0.46 |
| AHP Amplitude (mV) | -21.6±2.1 | -20.5±1.0 | 0.64 |
| AHP Duration (ms) | 210±42 | 192±31 | 0.73 |
| Input resistance (MΩ) | 175±19 | 178±14 | 0.90 |

**Figure S 2: Related to Figure 1.**
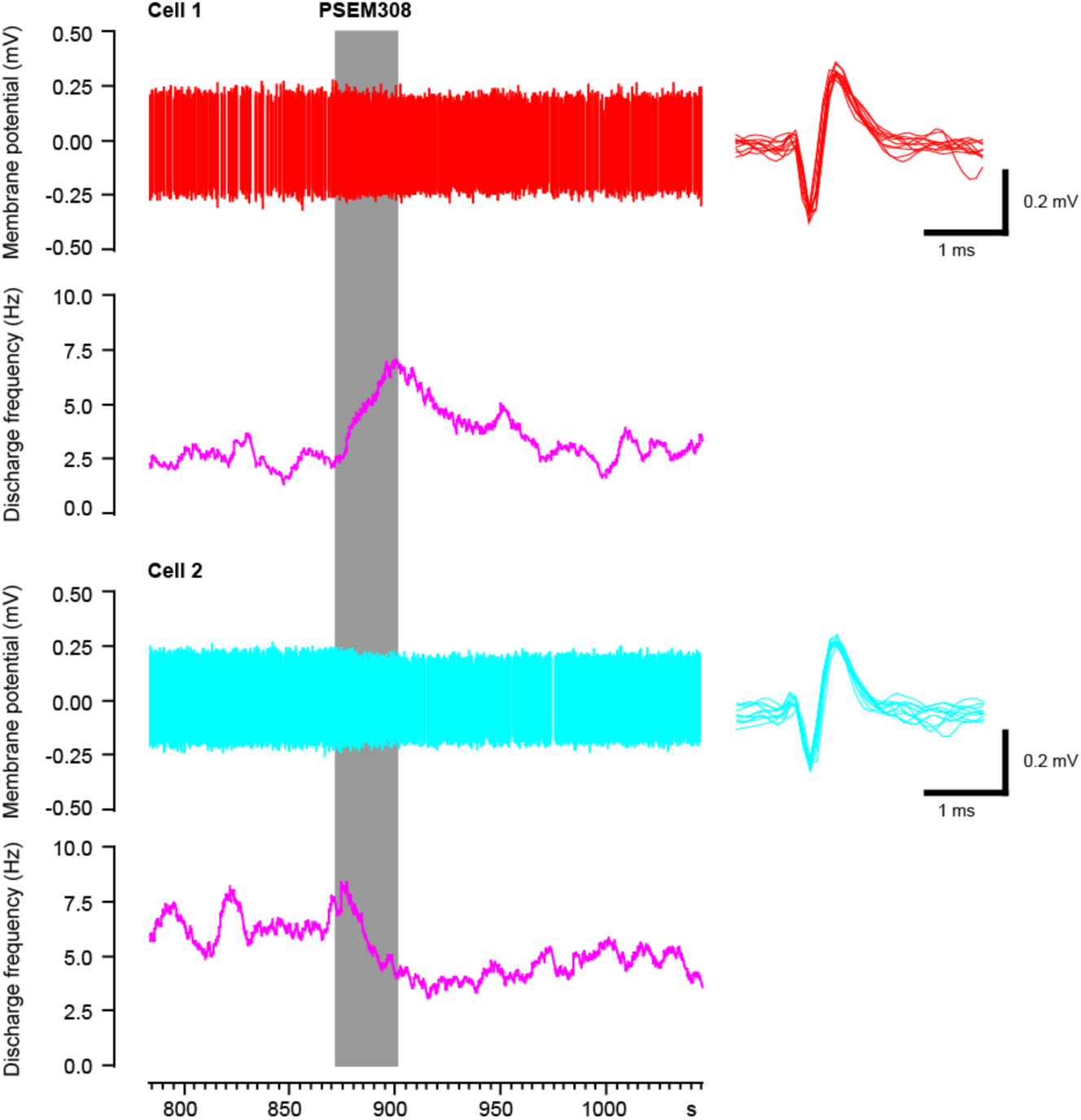
Simultaneous recording of one excited and one inhibited LC neuron in vivo with focal PSEM308 (1mM) pressure application.

**Figure S 3: Related to Figure 2.**
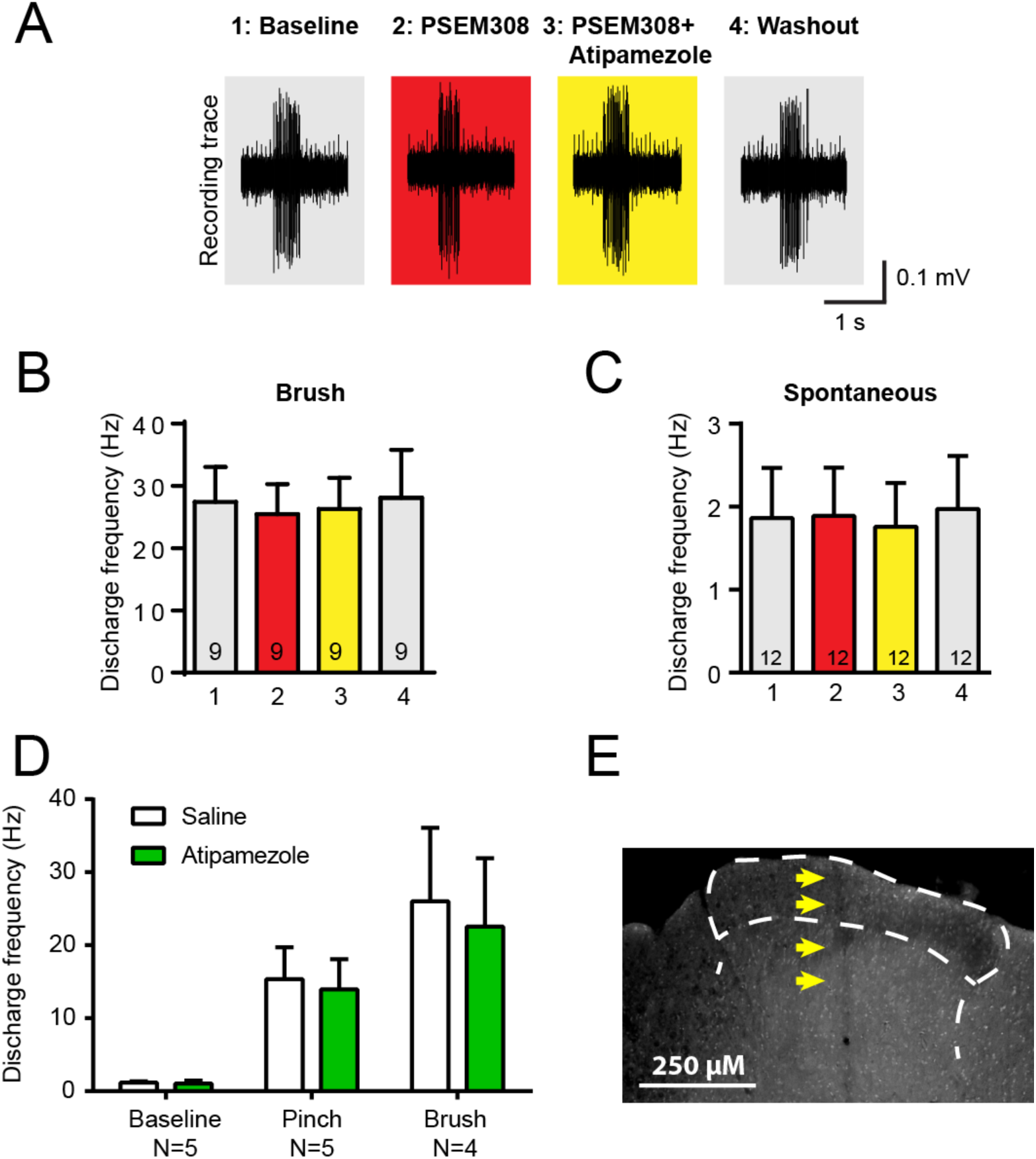
(A) Application of PSEM308 or atipamezole does not change spontaneous firing or brush evoked discharge of WDR in the dorsal horn Original traces of brush responses from WDR neurons during application of PSEM308 (10μM) or PSEM308 (10μM) + atipamezole (10μM) (B + C) Quantification of responses to brush and spontaneous firing (repeated measures ANOVA with Bonferroni’s multiple comparison p>0.05) (D) Quantification of neuronal responses to pinch, brush and spontaneous discharge frequency during saline or atipamezole application (paired t-test P>0.05) (E) Posthoc histology of the spinal dorsal horn outlining the electrode track Data are presented as mean ±SEM

**Figure S 4 Related to Figure 4.**
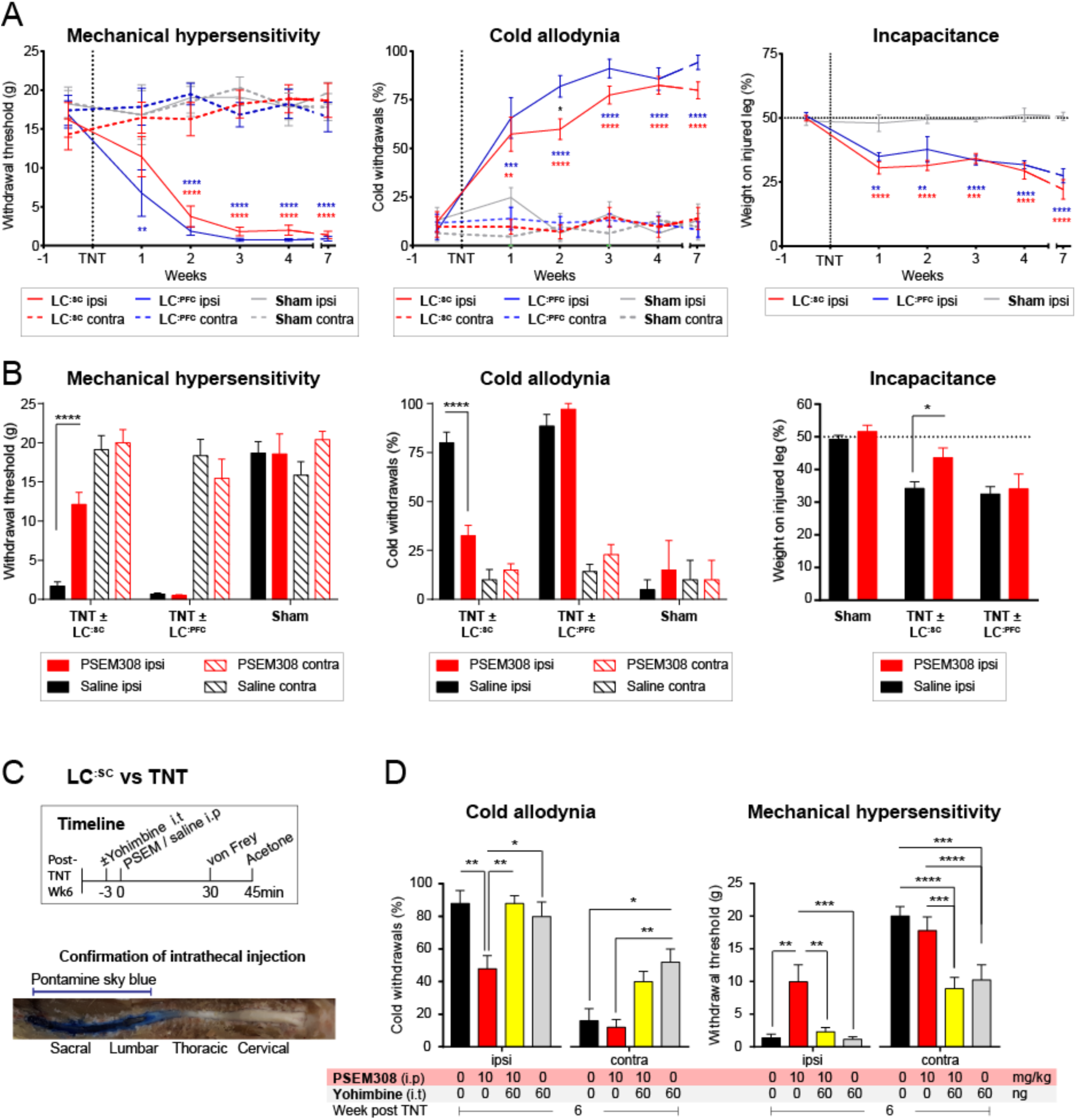
Development of neuropathic pain phenotype after tibial nerve transection and analgesic mechanism of chemogenetic activation of LC^:SC^ neurons. (A) Progression of mechanical sensitivity, cold sensitivity and incapacitance before and for 7 weeks after nerve injury (**LC^:SC^** N=8,**LC^:PFC^** N=7, sham N=6, 2-way repeated measures ANOVA with Bonferroni’s multiple comparison, * P<0.05, ** P<0.01, *** P<0.001, **** P<0.0001, red stars: **LC^:SC^** vs Sham, blue stars:**LC^:PFC^** vs Sham, black star **LC^:SC^** vs **LC^:PFC^**) (B) PSEM308 (10 mg/kg) alleviates ipsilateral hypersensitivity and improves incapacitance only in the **LC^:SC^** group 4 weeks after nerve injury (2-way repeated measures ANOVA with Bonferroni’s multiple comparison PSEM308 vs saline, * P<0.05, **** P<0.0001, **LC^:SC^** N=8,**LC^:PFC^** N=7, sham N=6) (C) Timeline of sensory testing and representative image of an intrathecal injection of pontamine sky blue 5 minutes before trans-cardiac perfusion (N=3). Dye was restricted to the caudal region of the spinal cord. (D) Ipsilateral chemogenetic analgesia was blocked by pre-treatment with yohimbine (repeated measures ANOVA with Bonferroni’s multiple comparison, * P<0.05, ** P<0.01, *** P<0.001). Note also that Intrathecal yohimbine (60ng i.t) lowered the mechanical and cold thresholds contralateral to nerve injury in the presence or absence of PSEM308 (10 mg/kg i.p) (2-way repeated measures ANOVA with Bonferroni’s multiple comparison, * P<0.05, ** P<0.01). Unmasking of contralateral sensitization was used as a positive control for successful intrathecal delivery of yohimbine (Hughes et al 2013).Data are presented as mean ±SEM

**Figure S 5:**
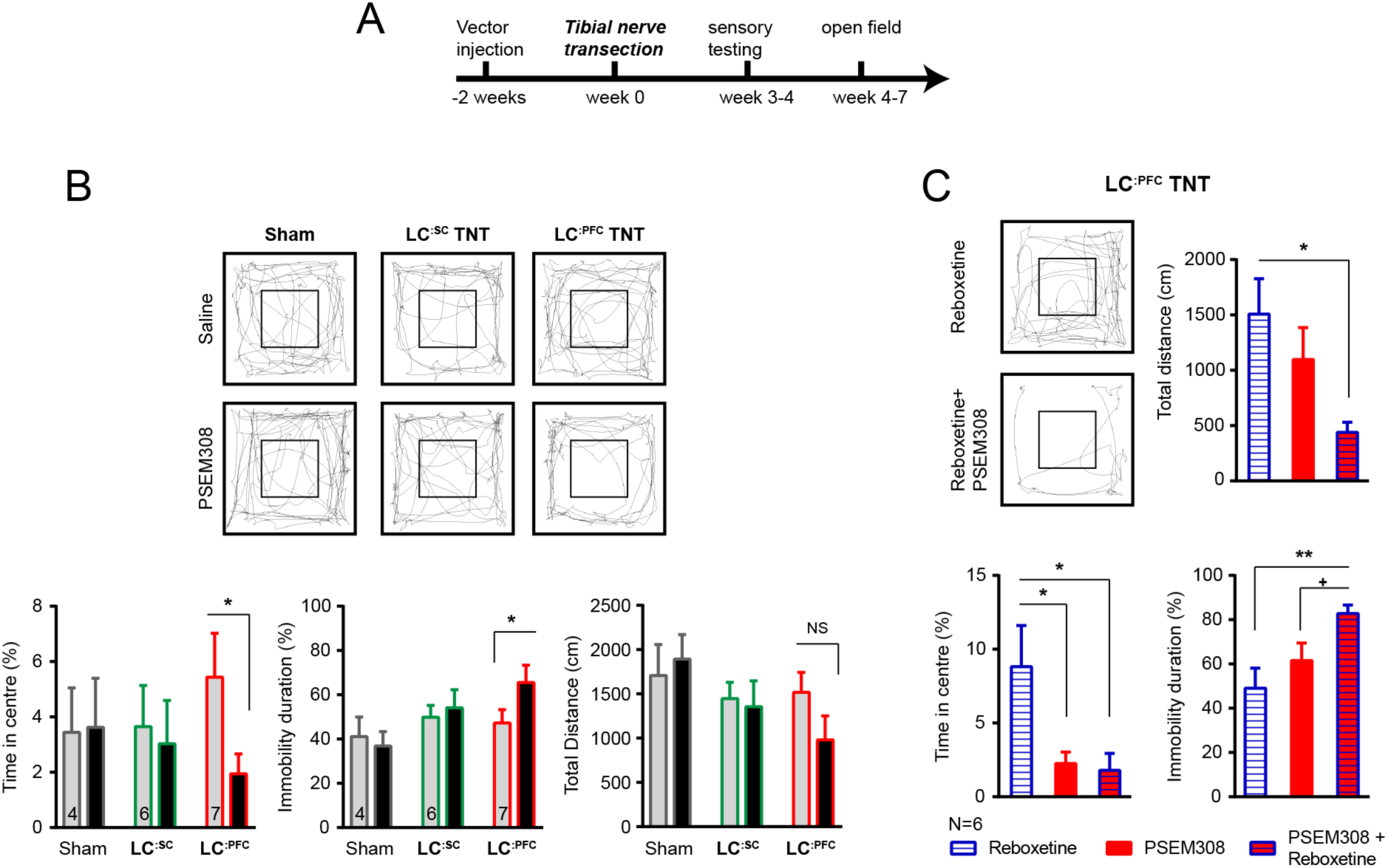
Chemogenetic stimulation of the LC^:PFC^ module is anxiogenic in chronic pain. (A) Timeline of nerve injury and open field testing (B) Representative heat maps showing activity in the open field test (OFT) after saline or PSEM308 (10 mg/kg) injection for sham, **LC^:SC^** and **LC^:PFC^** groups. Chemogenetic activation of **LC^:PFC^** reduced the time spent in the centre of the arena and increased immobility but PSEM308 had no effect on **LC^:SC^** transduced rats or sham rats (2-way repeated measures ANOVA with Bonferroni’s multiple comparison PSEM308 vs saline, * P<0.05). (C) Co-application of reboxetine (1 mg/kg) and PSEM308 (10 mg/kg) in **LC^:PFC^** rats increased immobility time, decreased the distance travelled and decreased the time spent in the centre of the arena as compared to reboxetine alone. Co-application of reboxetine and PSEM308 (10 mg/kg) also increased immobility as compared to PSEM308 alone (paired t-test PSEM308 vs PSEM308 + Reboxetine, + P<0.05).

